# DNA polymerase kappa supports repair-associated DNA synthesis and genome maintenance in postmitotic neurons

**DOI:** 10.64898/2026.10.01.756094

**Authors:** Gunjan Goyal, Premkumar Palanisamy, Ainesh Paul, Vijay L. Roy, Paolo Cifani, Anirban Paul, Shilpi Paul

## Abstract

Postmitotic neurons depend on continuous DNA repair to preserve genome integrity, yet the polymerases supporting repair-associated DNA synthesis remain poorly defined. Here, we identify DNA polymerase kappa (POLK), a Y-family translesion synthesis polymerase studied predominantly in proliferating cells, as a regulator of DNA synthesis and genome maintenance in postmitotic neurons. Using primary mouse cortical neurons and human induced pluripotent stem cell-derived forebrain neurons, we show that POLK localizes near newly synthesized DNA and repair-associated proteins, including ubiquitinated PCNA. POLK depletion reduces DNA synthesis in both mouse and human neurons and promotes nuclear and organelle remodeling, p21-associated senescence-like stress, inflammatory responses, and increased p38 MAPK immunoreactivity. In an Alzheimer’s disease-risk genetic background, POLK depletion also increases cytoplasmic double-stranded DNA and tau phosphorylation. Conversely, POLK overexpression enhances neuronal DNA synthesis while reducing DNA damage-associated signals, p38 MAPK immunoreactivity, cytoplasmic DNA, and Alzheimer’s disease-relevant phenotypes. These findings uncover a role for POLK in postmitotic neuronal genome maintenance, linking specialized DNA polymerase function to neuronal stress and vulnerability relevant to aging and neurodegeneration.

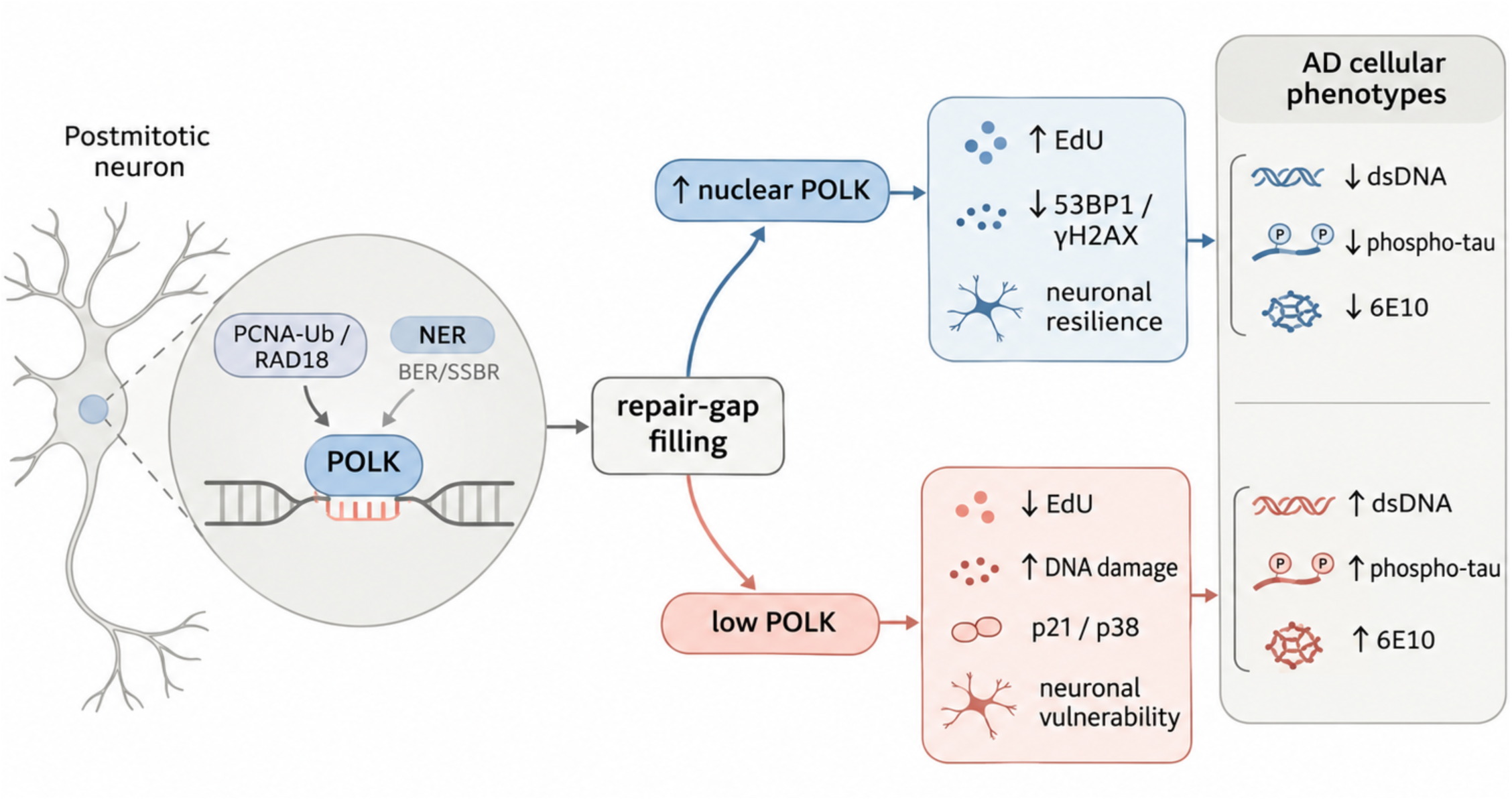

## INTRODUCTION

Neurons must preserve essentially the same genome for decades despite continuous endogenous DNA damage ^1–3^. Unlike proliferating cells, mature neurons are postmitotic and therefore rely heavily on continual DNA repair to preserve genome integrity throughout life. DNA repair frequently requires synthesis of new DNA to restore genome continuity. In base-excision repair (BER) and nucleotide-excision repair (NER), DNA polymerases fill gaps generated during lesion removal, whereas repair of DNA double-strand breaks by non-homologous end joining (NHEJ) can involve polymerase-mediated processing of DNA ends before ligation ^4^. Thus, incorporation of nucleotide analogs such as 5-ethynyl-2′-deoxyuridine (EdU) in postmitotic neurons can reveal nonreplicative DNA synthesis associated with repair. Two independent genome-wide studies have established that repair-associated DNA synthesis occurs extensively across the postmitotic neuronal genome. Repair-seq used EdU incorporation to map more than 60,000 reproducible repair hotspots in postmitotic human neurons, with repair enriched at active and evolutionarily constrained genomic regions ^5^. SAR-seq independently used EdU pulse to identify more than 55,000 sites of repair synthesis in human neurons and linked the strongest signals to active single-strand-break repair at neuronal enhancers ^6^. Together, these studies established that mature neurons undergo recurrent nonreplicative, repair-associated DNA synthesis ^7^. This continuous repair demand likely arises from both random endogenous DNA damage and programmed or physiological DNA breakage associated with transcription, topoisomerase activity, chromatin regulation, and active DNA demethylation ^2,3,8,9^. Recent work further indicates that completion of repair synthesis can itself become limiting in postmitotic neurons. Following cisplatin-induced DNA damage, NER can be efficiently initiated, yet low dNTP availability can impair gap filling, leaving persistent repair intermediates and promoting accumulation of single-and double-strand breaks and neuronal death ^10^. Thus, neuronal genome maintenance depends not only on lesion recognition and incision, but also on sufficient capacity to complete repair synthesis.

The polymerase repertoire that provides this capacity in mature neurons remains incompletely defined. POLb supports short-patch base-excision and single-strand-break repair, but neuronal repair synthesis involves additional pathways that may require other DNA polymerases. SAR-seq showed that impairment of POLb-dependent short-patch repair can shift synthesis toward longer repair tracts, indicating that additional polymerase systems contribute to neuronal gap filling ^6^. Which polymerases provide this broader repair-synthesis capacity in postmitotic neurons, particularly when synthesis must proceed across damaged or structurally challenging DNA templates, therefore remains unresolved.

DNA polymerase kappa (POLK) is a Y-family translesion synthesis (TLS) polymerase best known for lesion bypass during DNA replication, but TLS polymerases are increasingly recognized to have broader functions in DNA repair, gap filling, and genome maintenance outside conventional replication ^11^. POLK exemplifies this functional breadth. It contributes to nucleotide excision repair (NER), repair of DNA strand breaks, replication-independent repair of DNA interstrand crosslinks, and synthesis across damaged or structurally challenging DNA templates ^12–15^. Importantly, POLK function is not restricted to S phase. POLK participates in replication-independent interstrand-crosslink repair in noncycling cells, demonstrating that its genome-maintenance functions can operate independently of genome replication ^13^. Conversely, in proliferating cells, POLK promotes DNA synthesis and replication-fork recovery following hydroxyurea-induced nucleotide deprivation ^16^. During human NER, POLK and DNA polymerase delta (POLδ) contribute to repair synthesis through distinct recruitment mechanisms, with POLK engagement promoted by ubiquitinated PCNA (PCNA-Ub) and XRCC1. RAD18-dependent PCNA ubiquitination provides an established mechanism for POLK recruitment to damaged DNA ^12,17^. Our recent work further extends these noncanonical functions to the nucleolus, where POLK associates with ribosomal DNA (rDNA) and contributes to DNA synthesis and recovery of rRNA production following nucleolar genotoxic stress ^18^.

Although Y-family TLS polymerases are often described as error-prone, POLK fidelity depends strongly on lesion and sequence context. POLK can accurately bypass several N2-guanine adducts, including bulky benzo[a]pyrene diol epoxide-N2-deoxyguanosine, while POLK loss increases mutagenesis associated with N2-carboxyalkyl-2′-deoxyguanosine lesions. Recent somatic mutation profiling further demonstrated that POLK suppresses mutations arising from endogenous guanine damage in mouse tissues ^19–22^. Thus, POLK can protect genome integrity rather than being intrinsically mutagenic. Genome-wide studies in proliferating human cells revealed preferential POLK activity in early-replicating genomic regions, GC-rich sequences, accessible chromatin, and gene promoters, while other studies implicated POLK in maintaining genome stability at G-rich and structurally challenging DNA sequences ^15,23,24^.

Genetic studies further support an in vivo genome-protective role for POLK. Constitutive germline *Polk*-null mice develop an age-dependent spontaneous mutator phenotype and reduced survival ^25^, while *Polk* loss in an *Apoe*-null hypercholesterolemic background increases mutation burden, including in the brain, and shortens survival under high-cholesterol conditions ^26^. Although these systemic germline models demonstrate a role for POLK in genome protection, they do not establish its cell-autonomous function in mature central nervous system (CNS) neurons. Previous neuronal evidence comes from peripheral dorsal root ganglion neurons, in which POLK depletion reduced cisplatin-induced DNA repair synthesis ^27^. Thus, whether POLK contributes to endogenous repair-associated DNA synthesis and genome maintenance in postmitotic CNS neurons remains unresolved.

Our previous investigation of the aging mouse brain identified POLK as a prominent nuclear protein in postmitotic CNS neurons, with higher nuclear abundance in GABAergic interneurons than in excitatory pyramidal neurons. Nuclear POLK also localized near DNA damage markers and repair-associated proteins. With aging, nuclear POLK progressively declined while cytoplasmic POLK accumulated, coinciding with increased neuronal DNA damage-associated signals. Cytoplasmic POLK in aged neurons was associated with stress granules and endosomal/lysosomal compartments, whereas increased neuronal activity promoted its nuclear localization, suggesting that POLK distribution is dynamically regulated rather than simply diminished with age ^28^. Together, these observations raised the possibility that age-associated loss of nuclear POLK compromises neuronal genome maintenance. However, whether POLK contributes to repair-associated DNA synthesis in postmitotic CNS neurons and how changes in its abundance affect neuronal function remained unknown.

Persistent genome damage can also drive broader remodeling of postmitotic neuronal state. Aged neurons with sustained DNA damage responses develop p21- and p38-associated senescence-like phenotypes accompanied by inflammatory and organelle stress ^29^. Persistent DNA damage and accumulation of cytoplasmic double-stranded DNA (dsDNA) can additionally engage cGAS–STING innate immune signaling, which has been linked to aging-associated neuroinflammation, neurodegeneration, and senescence-like states in postmitotic neurons ^30,31^. These pathways are particularly relevant to Alzheimer disease (AD), in which p38α/MAPK14 signaling, cytoplasmic DNA sensing, and persistent neuronal stress have been associated with amyloid- and tau-related pathology ^32,33^. Together, these observations raise the possibility that declining neuronal repair capacity could couple persistent genome damage to inflammatory and stress states that increase vulnerability to AD-associated pathology.

Here, we use primary mouse cortical neurons and human induced pluripotent stem cell (hiPSC) derived forebrain neurons generated through developmental differentiation to define the consequences of POLK loss and gain of function. We test whether POLK localizes near newly synthesized DNA, whether POLK abundance influences neuronal EdU incorporation across mouse and human neuronal models, which DNA repair proteins and molecular contexts are associated with neuronal POLK, and whether altering POLK levels modifies genome damage, persistent stress, and AD-relevant cellular phenotypes. Our findings implicate POLK in repair-associated DNA synthesis in mature neurons and support a model in which POLK abundance influences neuronal genome maintenance, persistent stress, and AD-relevant cellular phenotypes.

## RESULTS

### POLK depletion reduces neuronal complexity and remodels nuclear architecture

To define the consequences of reducing POLK in postmitotic neurons, we established complementary mouse and human neuronal systems (Fig. 1A). Primary cortical neurons were isolated from P0–P1 mice, enriched by magnetic sorting, matured to DIV10, and treated with previously validated siRNA or lentiviral shRNA targeting *Polk* before analysis at DIV13 (Fig. 1Ai; ^28^). Both approaches significantly reduced *Polk* mRNA (Fig. 1Bi,ii).

**Figure 1.**
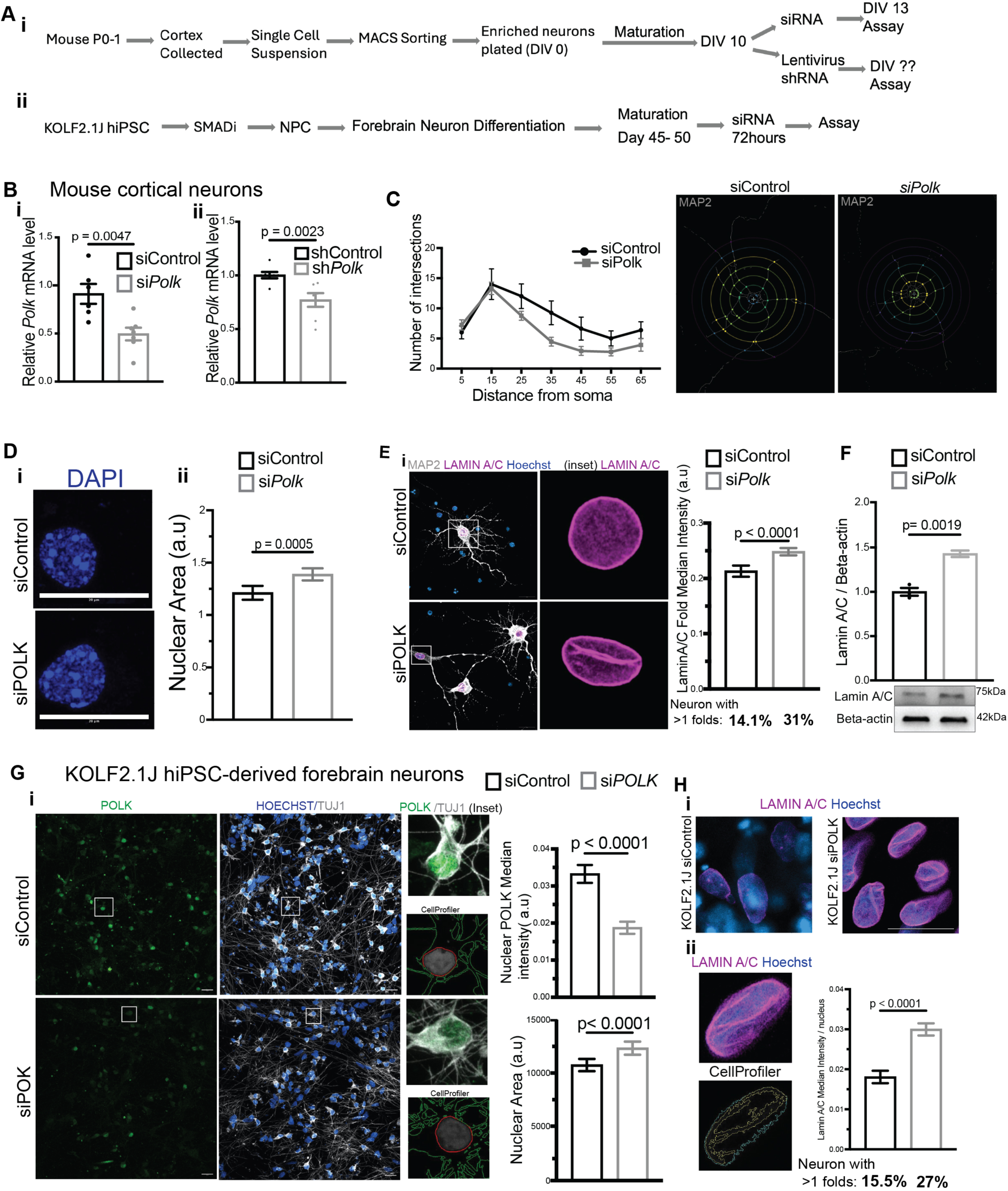
POLK depletion reduces neuronal complexity and remodels nuclear architecture in mouse and human neurons. (A) Experimental paradigms. (i) Primary cortical neurons were isolated from P0-P1 mice, dissociated, enriched by MACS, plated at DIV0, matured to DIV10, treated with siRNA or lentiviral shRNA targeting Polk, and analyzed at DIV13. (ii) KOLF2.1J hiPSCs were differentiated through an NPC intermediate into forebrain neurons, matured for 45-50 days, treated with siRNA for 72 h, and analyzed. (B) Reduction of Polk mRNA following (i) siRNA (p = 0.0047) or (ii) shRNA (p = 0.0023) treatment in mouse cortical neurons. (C) Sholl analysis and representative MAP2-positive mouse cortical neurons showing reduced neurite intersections following POLK depletion. (D) Nuclear morphology in mouse cortical neurons. (i) Representative DAPI images. (ii) Increased nuclear area following POLK depletion (p = 0.0005). (E) Representative MAP2, lamin A/C, and Hoechst images and quantification of nuclear lamin A/C intensity. POLK depletion increased lamin A/C intensity (P < 0.0001) and the proportion of neurons containing >1 nuclear fold from 14.1% to 31.0%. (F) Immunoblot and quantification of lamin A/C relative to β-actin (p = 0.0019). (G) POLK depletion in KOLF2.1J-derived forebrain neurons. Representative POLK, TUJ1, and Hoechst images with CellProfiler segmentation and quantification showing reduced nuclear POLK and increased nuclear area (both P < 0.0001). (H) Representative lamin A/C/Hoechst images and CellProfiler-based analysis showing increased nuclear lamin A/C intensity (P < 0.0001); neurons with >1 nuclear fold increased from 15.5% to 27.0%.

In parallel, KOLF2.1J hiPSCs, a deeply characterized reference line for disease modeling ^34^, were differentiated through a neural progenitor intermediate into forebrain neurons and matured for 45–50 days before 72-h treatment with pooled POLK-targeting or non-targeting control siRNAs (Fig. 1Aii). Immunostaining confirmed that MAP2-positive neurons lacked detectable Ki67 expression, whereas the few Ki67-positive nuclei were restricted to MAP2-negative cells, supporting the postmitotic identity of the differentiated neuronal population (Supplementary Fig. S1B). Although structural variants have been characterized in KOLF2.1J, control and POLK-depleted neurons were both generated from the same parental background and differentiated in parallel, thus holding constitutive genetic variation constant across experimental conditions ^35,36^. Because our prior *in vivo* work identified neuronal-class differences in nuclear POLK, we used developmental forebrain differentiation rather than restricting the study to a single transcription factor programmed neuronal identity.

AD provides a particularly relevant disease context for investigating POLK because aging, its strongest risk factor, is accompanied by accumulating neuronal DNA damage and declining genome-maintenance capacity, processes increasingly implicated in AD pathogenesis ^1–3^. Moreover, APOE4, the strongest common genetic risk factor for late-onset AD, is associated with altered neuronal lipid homeostasis, APP processing, tau phosphorylation, and cellular stress ^37–40^. We therefore incorporated a CRISPR-engineered homozygous APOE e4/e4 hiPSC line generated from the KOLF2.1J parental background by The Jackson Laboratory to examine the consequences of altering POLK abundance in an AD-risk genetic context. The shared parental background enabled us to extend the investigation of POLK-dependent neuronal genome maintenance to a complementary genetic model relevant to AD. Consistent with this rationale, single-nucleus RNA sequencing of matched-stage KOLF2.1J and APOE e4/e4 neuronal cultures identified glutamatergic and GABAergic populations together with additional neuronal transcriptional states (Supplementary Fig. S1C). Partial POLK transcript reduction in KOLF2.1J and APOE e4/e4 -derived neurons was independently verified by qPCR (Fig. 3E).

POLK knockdown altered neuronal morphology and nuclear architecture. Sholl analysis of MAP2-positive mouse cortical neurons showed fewer neurite intersections with increasing distance from the soma after Polk depletion, consistent with reduced arbor complexity (Fig. 1C). si*Polk*-treated mouse neurons also displayed enlarged nuclei, with a significant increase in nuclear area (p = 0.0005; Fig. 1Di,ii). Lamin A/C immunostaining showed increased nuclear intensity and more prominent nuclear-envelope folding; the fraction of neurons displaying more than one fold increased from 14.1% to 31.0% after POLK depletion (Fig. 1E). Immunoblotting independently confirmed increased lamin A/C abundance (p = 0.0019; Fig. 1F).

Similar nuclear phenotypes were observed in hiPSC-derived forebrain neurons. POLK depletion in KOLF2.1J-derived neurons reduced nuclear POLK and increased nuclear area (both P < 0.0001; Fig. 1G). Lamin A/C intensity was increased, and the fraction of neurons displaying more than one nuclear-envelope fold increased from 15.5% to 27.0% (Fig. 1H). Thus, partial POLK loss reduces neuronal arbor complexity in mouse cortical neurons and produces a reproducible nuclear-lamina remodeling phenotype across mouse and human neuronal systems.

### POLK depletion induces a multidimensional senescence-like neuronal stress state with lysosomal and mitochondrial remodeling

We next asked whether the nuclear remodeling following POLK knockdown was accompanied by a broader persistent stress state. Mouse cortical neurons with reduced POLK exhibited increased senescence-associated β-galactosidase (SA-β-gal) activity relative to controls (Fig. 2Ai-iii). Because mature neurons are already postmitotic and SA-β-gal reflects increased lysosomal activity rather than a neuron-specific senescence state, neither cell-cycle arrest nor SA-β-gal alone is sufficient to define neuronal senescence. We therefore interpreted this phenotype together with nuclear morphology, organelle homeostasis, stress signaling, and inflammatory changes, consistent with multidimensional criteria proposed for neuronal senescence or ’neurescence’ ^41^.

**Figure 2.**
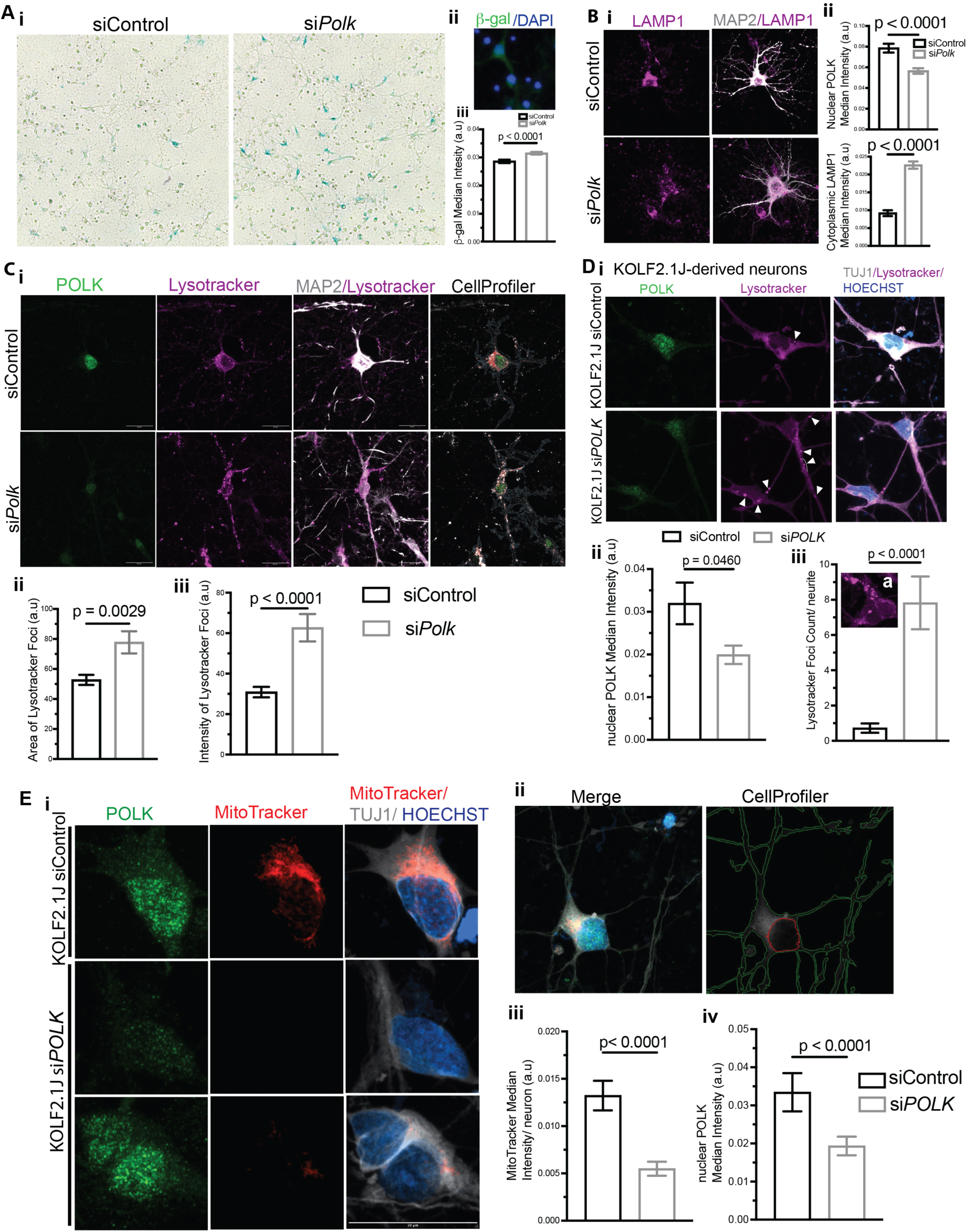
POLK depletion induces senescence-associated and organelle-remodeling phenotypes in neurons. (A) Senescence-associated β-galactosidase after POLK depletion. (i) Representative siControl and siPolk cultures. (ii) Representative β-gal/DAPI image. (iii) Quantification showing increased β-gal signal (P < 0.0001). (B) LAMP1 staining in mouse cortical neurons. (i) Representative LAMP1 and MAP2 images. (ii) POLK depletion reduced nuclear POLK and increased cytoplasmic LAMP1 intensity (both P < 0.0001). (C) LysoTracker analysis in mouse cortical neurons. (i) Representative POLK, LysoTracker, MAP2, and CellProfiler images. Quantification shows increased (ii) LysoTracker-positive area (p = 0.0029) and (iii) LysoTracker intensity (P < 0.0001). (D) LysoTracker analysis in KOLF2.1J-derived forebrain neurons. (i) Representative POLK, LysoTracker, TUJ1, and Hoechst images; arrowheads mark LysoTracker-positive structures along neurites. (ii) Reduced nuclear POLK (p = 0.0460) and (iii) increased LysoTracker-positive puncta per neurite (P < 0.0001) following POLK depletion. (E) MitoTracker analysis in human forebrain neurons. (i) Representative POLK, MitoTracker, TUJ1, and Hoechst images. (ii) Representative merged image and CellProfiler segmentation. Quantification shows (iii) reduced MitoTracker intensity and (iv) reduced nuclear POLK following POLK depletion (both P < 0.0001).

POLK depletion also altered lysosomal organization in mouse cortical neurons, as indicated by increased cytoplasmic LAMP1 staining accompanying reduced nuclear POLK (Fig. 2Bi,ii). LysoTracker imaging similarly showed increased acidic-organelle-associated signal, with greater LysoTracker-positive area and fluorescence intensity after POLK depletion (Fig. 2Ci-iii). A comparable phenotype was observed in KOLF2.1J-derived human forebrain neurons, where reduced nuclear POLK was associated with a marked increase in LysoTracker-positive puncta along neuronal processes (Fig. 2Di-iii).

POLK reduction also altered mitochondrial-associated staining. In human forebrain neurons, siPOLK treatment significantly reduced MitoTracker fluorescence together with the expected reduction in nuclear POLK (Fig. 2Ei-iv). Lysosomal and mitochondrial remodeling are recurring features of neuronal senescence-like states produced in experimental models including long-term cultured cortical neurons with impaired autophagy and p21-associated postmitotic neuronal senescence ^42,43^. Our findings indicate coordinated organelle remodeling following POLK depletion.

### POLK deficiency engages inflammatory and senescence-associated stress programs

We next examined whether the structural and organelle phenotypes associated with POLK reduction were accompanied by inflammatory and senescence-associated stress signaling. In mouse cortical neurons, si*Polk* altered expression of several stress-associated transcripts, including *Cdkn1a*, *Cdkn2a/p16Ink4a*, *Cdkn2a/p19Arf*, *Ccl2*, *Cxcl1*, and *Il6*, with *Cxcl1* significantly increased (p = 0.0417; Fig. 3Ai). Independent shRNA-mediated POLK reduction significantly increased *Ccl2* (p = 0.0365; Fig. 3Aii), supporting an inflammatory response across independent knockdown approaches. CCL2/MCP-1 is also increased and secreted in senescence-like cortical-neuron cultures ^42^. At the protein level, POLK depletion significantly increased CDKN1A/p21 abundance (p = 0.0433; Fig. 3B), consistent with previous studies linking persistent DNA damage signaling to p21-dependent senescence-like responses in postmitotic neurons ^44,45^.

**Figure 3.**
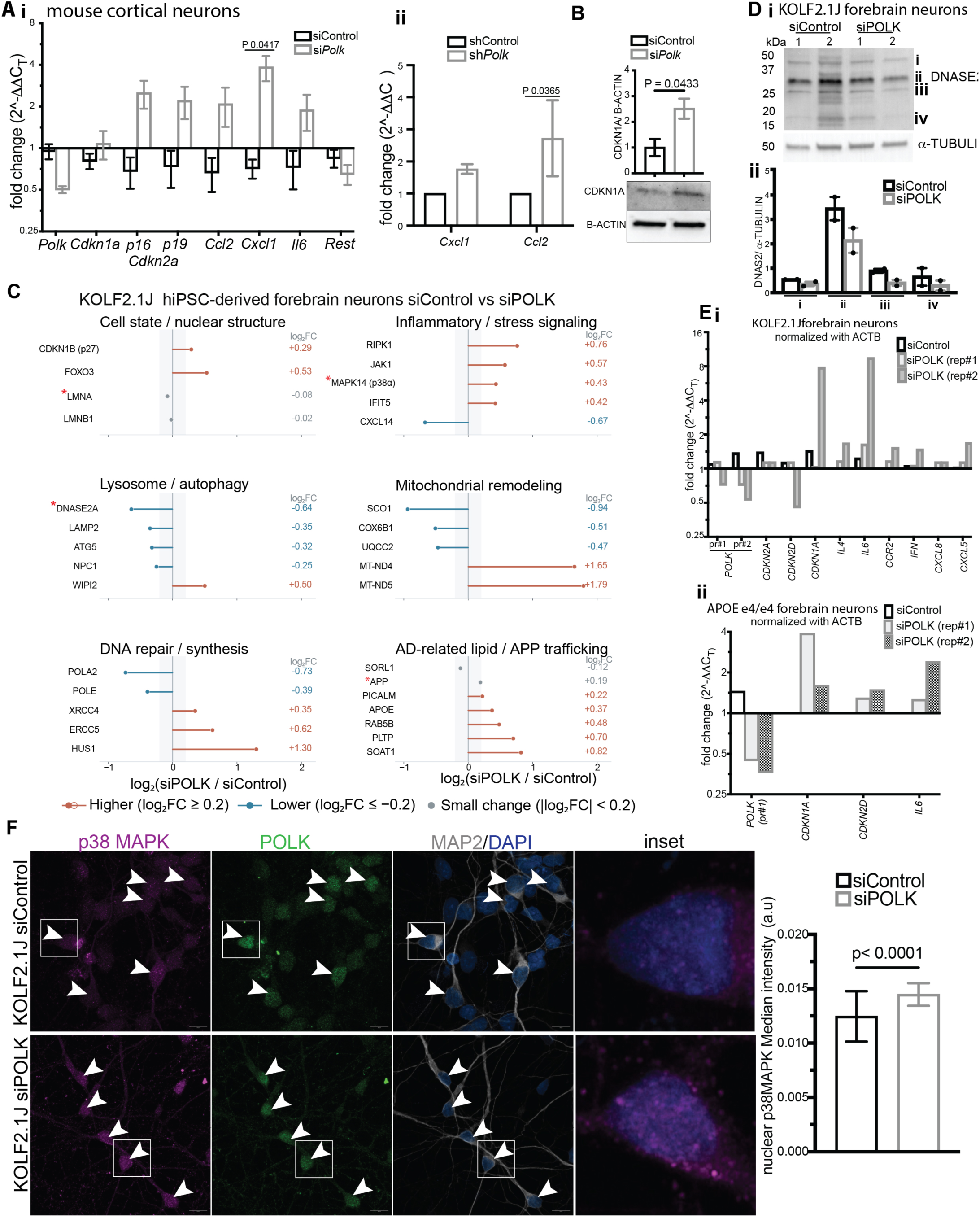
POLK depletion engages inflammatory and senescence-associated stress programs. (A) qPCR analysis in mouse cortical neurons. (i) Expression of Polk, Cdkn1a, Cdkn2a/p16Ink4a, Cdkn2a/p19Arf, Ccl2, Cxcl1, Il6, and Rest after siRNA-mediated POLK depletion; Cxcl1 was significantly increased (p = 0.0417). (ii) Independent shRNA depletion increased Ccl2 (p = 0.0365). (B) Immunoblot and quantification of CDKN1A/p21 relative to β-actin after POLK depletion (p = 0.0433). (C) Exploratory proteomic profiling of wild-type hiPSC-derived forebrain neurons treated with control or POLK siRNA. One sample per condition was analyzed; fold changes are displayed descriptively for selected proteins grouped into nuclear/cell state, inflammatory/stress signaling, lysosome/autophagy, mitochondrial remodeling, DNA repair/synthesis, and AD-related lipid/APP-trafficking pathways. (D) DNASE2 protein in wild-type human forebrain neurons. (i) Immunoblot showing multiple DNASE2-reactive molecular species in two independent control and siPOLK neuronal preparations. Bands i-iv are indicated according to apparent molecular weight and were quantified individually; they were not assigned to specific precursor or processed states. Alpha-tubulin served as loading control. (ii) Densitometric analysis of each indicated DNASE2-reactive band normalized to α-tubulin; with two preparations per condition, these measurements are interpreted descriptively. (E) qPCR of stress-and inflammatory-response genes after POLK depletion in (i) wild-type and (ii) APOE e4/e4 forebrain neurons. The human p19 assay corresponds to CDKN2D/p19INK4d, distinct from mouse Cdkn2a/p19Arf. Independent siPOLK preparations are shown relative to control and normalized to ACTB. (F) Representative p38 MAPK, POLK, MAP2, and DAPI images in wild-type human forebrain neurons and quantification showing increased nuclear p38 MAPK immunoreactivity after POLK depletion (p < 0.0001).

To identify broader pathways associated with POLK reduction in human neurons, we performed exploratory proteomic profiling of wild-type hiPSC-derived forebrain neurons treated with control or POLK siRNA (Supplementary Table 1). Because this analysis contained one sample per condition, fold changes were used for pathway nomination rather than statistical inference. POLK depletion was associated with changes in proteins related to nuclear and cell-state regulation, inflammatory and stress signaling, lysosomal and autophagy-related processes, mitochondrial function, DNA repair and synthesis, and AD-related lipid and APP trafficking (Fig. 3C). Notable changes included increased MAPK14/p38α, RIPK1, JAK1, and IFIT5; reduced DNASE2A, LAMP2, and ATG5; changes in mitochondrial proteins; and altered abundance of DNA-repair and synthesis factors including POLA2, POLE, XRCC4, ERCC5, and HUS1. This pattern additionally suggested remodeling of innate immune and interferon-associated signaling following reduction of POLK. These exploratory data were used to nominate proteins and pathways for independent examination rather than to infer pathway activity.

Among the proteins reduced in the exploratory proteome, DNASE2A was of particular interest because of its role in intracellular DNA clearance. Immunoblotting of cytoplasmic fractions from independent human forebrain-neuron preparations detected multiple DNASE2-reactive molecular species spanning approximately 20–45 kDa, several of which were reduced following POLK depletion (Fig. 3Di,ii), providing orthogonal support for the proteomic observation. The DNASE2-reactive bands were quantified separately without assigning specific molecular identities or processing states.

Independent qPCR analyses in KOLF2.1J-derived forebrain neurons confirmed partial POLK transcript reduction and showed recurrent increases in stress- and inflammatory-response genes, most prominently CDKN1A and IL6, together with changes in additional cytokine and chemokine transcripts (Fig. 3Ei). Matched-stage APOE e4/e4 forebrain neurons showed similar increases in CDKN1A and IL6 following POLK knockdown (Fig. 3Eii). CDKN2D/p19INK4d, previously associated with neuronal senescence and tau neuropathology ^46^, showed a less consistent response. Thus, p21 and inflammatory signaling were the more consistent transcriptional responses to POLK depletion across both human neuronal backgrounds.

Finally, MAPK14/p38a was prioritized for orthogonal validation because it emerged as an increased candidate in the exploratory proteome and has been linked independently to DNA-damage-associated neuronal senescence and AD. POLK depletion significantly increased nuclear p38α MAPK immunoreactivity in MAP2-positive human forebrain neurons (P < 0.0001; Fig. 3F). This finding is consistent with p38 MAPK-associated signaling in aged postmitotic neurons exhibiting DNA damage-driven, p21-dependent senescence-like phenotypes ^44^. Its relevance to AD is further supported by integrative analyses identifying the p38 MAPK–MK2 axis as a shared upregulated pathway in human AD brain tissue and three-dimensional human AD models ^33^. Together with the SA-β-gal activity, nuclear and lysosomal remodeling, and mitochondrial-associated changes in Fig. 2, the induction of p21 and inflammatory signaling together with increased nuclear p38a MAPK supports a convergent, multidimensional senescence-like neuronal stress state following POLK reduction.

### POLK localizes near sites of ongoing DNA synthesis in postmitotic neurons

The nuclear remodeling, inflammatory signaling, and senescence-like stress observed following POLK depletion raised the possibility of impaired neuronal genome maintenance. To investigate a potential underlying mechanism, we examined whether POLK localizes near sites of ongoing DNA synthesis in postmitotic neurons. Building on previous Repair-seq and SAR-seq studies demonstrating nonreplicative DNA repair synthesis in mature human neurons ^6,47^, we adopted the EdU-labeling conditions used in SAR-seq (20 μM for 18 h) to examine POLK proximity to newly synthesized DNA.

We adapted a single-cell in situ protein proximity assay for nascent DNA (SIRF) in mature KOLF2.1J-derived forebrain neurons. Following EdU incorporation and biotin conjugation by click chemistry, POLK proximity to biotin-labeled newly synthesized DNA was assessed by proximity ligation assay (PLA) using anti-POLK and anti-biotin antibodies (Fig. 4A). POLK-biotin-EdU SIRF produced sparse, discrete PLA puncta in MAP2-positive neurons, with typically one or a few puncta detected within the nucleus and occasional additional puncta in the neuronal soma, whereas the negative-control condition showed little or no PLA signal (Fig. 4Bi–iii). The focal nuclear PLA pattern suggests that POLK localizes near discrete sites of ongoing DNA synthesis. To further examine this spatial relationship, we analyzed endogenous POLK and EdU incorporation in primary mouse cortical neurons using an independent imaging approach. MAP2-positive neurons contained nuclear EdU signal that spatially overlapped with endogenous POLK immunoreactivity (Fig. 4Ci). Puncta level analyses across individual neurons showed positive relationships between POLK and EdU intensity in most cells analyzed (Fig. 4Cii). Thus, complementary proximity and spatial-imaging approaches in human and mouse neurons independently position POLK near sites of ongoing neuronal DNA synthesis.

**Figure 4.**
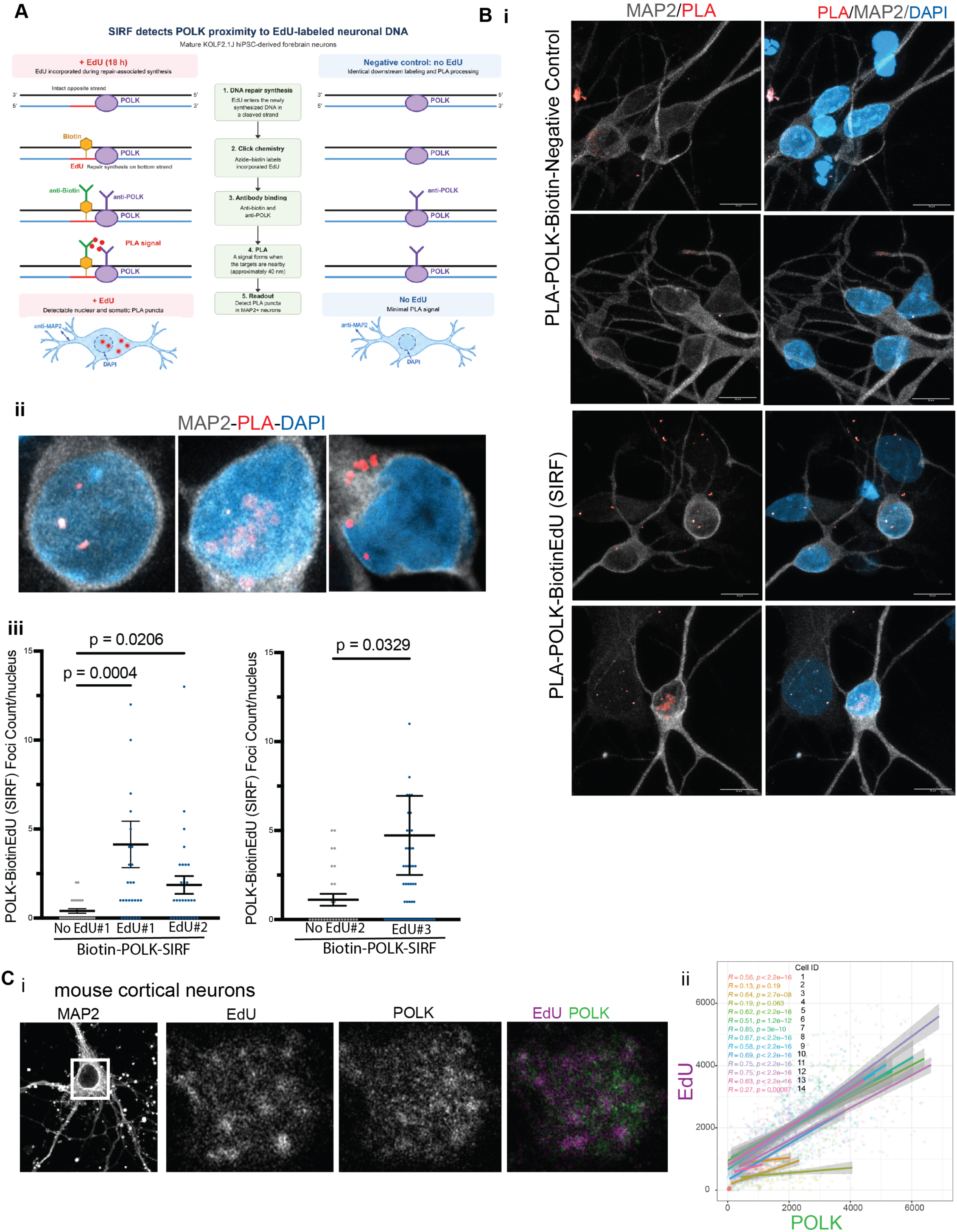
POLK localizes near sites of ongoing DNA synthesis in postmitotic neurons. (A) Schematic of the POLK–biotin–EdU SIRF assay. Mature KOLF2.1J-derived forebrain neurons were exposed to 20 μM EdU for 18 h. EdU incorporation was followed by azide–biotin click chemistry, anti-biotin and anti-POLK antibody incubation, and proximity ligation assay (PLA). Cells without EdU were subjected to the same downstream processing as negative controls. The schematic depicts repair-associated synthesis at a cleaved DNA strand with the opposite strand intact and is conceptual. (B) POLK–biotin–EdU SIRF in human forebrain neurons. (i) Representative MAP2, PLA, and DAPI images comparing negative-control and SIRF conditions. (ii) Enlarged nuclear examples showing discrete SIRF puncta. (iii) Nuclear SIRF puncta per neuron in independently labeled conditions with the indicated pairwise P values (P = 0.0004, P = 0.0206, and P = 0.0329). (C) Endogenous POLK and EdU signal in mouse cortical neurons. (i) Representative MAP2, EdU, POLK, and merged images. (ii) Relationships between POLK and EdU fluorescence intensities for individual neurons; cell-wise correlations and P values are indicated.

### POLK depletion reduces DNA synthesis in mouse cortical and human forebrain neurons

Following our observation that POLK localizes near sites of ongoing DNA synthesis, we next investigated whether reducing POLK abundance alters DNA synthesis in mature human forebrain neurons. KOLF2.1J-derived forebrain neurons were treated with POLK siRNA for 48 h and exposed to 20 μM EdU for the final 18 h (Fig. 5A). EdU-positive nuclei were detected in TUJ1-positive neurons lacking detectable GFAP immunoreactivity, distinguishing neuronal DNA synthesis from that occurring in GFAP-positive astroglial cells within differentiated cultures (Supplementary Fig. S2A). POLK depletion significantly reduced neuronal EdU intensity (p = 0.0002; Fig. 5Bi,ii), alongside reduced nuclear POLK (P < 0.0001; Fig. 5Biii), indicating decreased ongoing DNA synthesis in mature human forebrain neurons.

**Figure 5.**
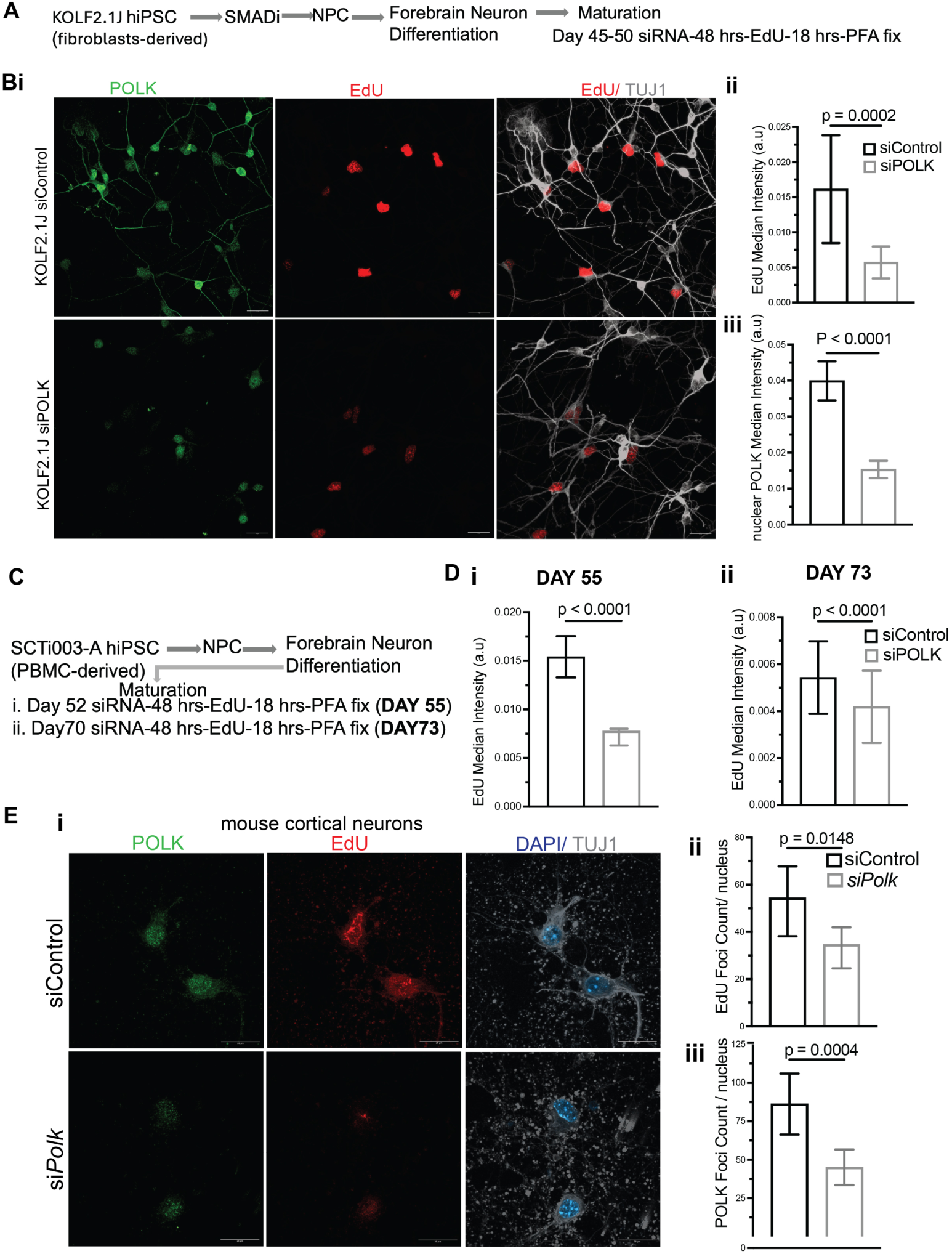
POLK depletion reduces DNA synthesis in human forebrain and primary mouse cortical neurons. (A) Experimental workflow for KOLF2.1J hiPSC-derived forebrain neurons, including forebrain differentiation, maturation to days 45–50, 48-h siRNA treatment, and 20 μM EdU for the final 18 h before fixation. (B) (i) Representative POLK, EdU, and TUJ1 images comparing siControl and siPOLK neurons. (ii) Quantification of neuronal EdU median intensity (p = 0.0002) and (iii) nuclear POLK median intensity (P < 0.0001). (C) Differentiation and EdU labeling workflow for independent SCTI003-A-derived forebrain neurons analyzed at days 55 and 73. (D) Reduced EdU median intensity after POLK depletion at (i) day 55 and (ii) day 73 (both P < 0.0001); additional day-73 images and nuclear POLK quantification are shown in Supplementary Fig. 2. (E) Primary mouse cortical neurons. (i) Representative POLK, EdU, and DAPI/TUJ1 images comparing siControl and siPolk. Quantification of (ii) EdU-positive foci per nucleus (p = 0.0148) and (iii) POLK-positive foci per nucleus (p = 0.0004).

To assess reproducibility across independent human hiPSC lines, we repeated the analysis in SCTI003-A-derived forebrain neurons, generated from a different donor using a distinct reprogramming method, at differentiation days 55 and 73 (Fig. 5C). POLK depletion significantly reduced neuronal EdU incorporation at day 55 (p < 0.0001; Fig. 5Di), with a similar reduction observed at day 73 (p < 0.0001; Fig. 5Dii). Representative imaging and nuclear POLK quantification at day 73 are presented in Supplementary Fig. 2B. Together with the findings in KOLF2.1J-derived neurons, these results demonstrate that reduced POLK abundance is associated with decreased neuronal DNA synthesis across two independent donor-derived hiPSC lines and at different stages of neuronal maturation.

To determine whether this effect extends to primary CNS neurons, we examined EdU incorporation in mouse cortical neurons after siRNA-mediated *Polk* depletion (Fig. 5Ei). Consistent with the results in human forebrain neurons, si*Polk* treatment reduced the number of EdU-positive foci per nucleus (p = 0.0148; Fig. 5Eii), accompanied by a reduction in POLK-positive foci per nucleus (p = 0.0004; Fig. 5Eiii). These findings extend the POLK depletion-associated reduction in DNA synthesis across mouse and human neuronal models.

### POLK occupies multiple DNA-repair environments associated with neuronal repair synthesis

To identify candidate proteins associated with nuclear POLK in human neurons, we performed exploratory nuclear POLK immunoprecipitation followed by mass spectrometry (IP–MS) in KOLF2.1J-derived forebrain neurons, using an IgG immunoprecipitation as control (Fig. 6A, Supplementary Table 2). Successful POLK immunoprecipitation was independently confirmed by immunoblotting, although

**Figure 6.**
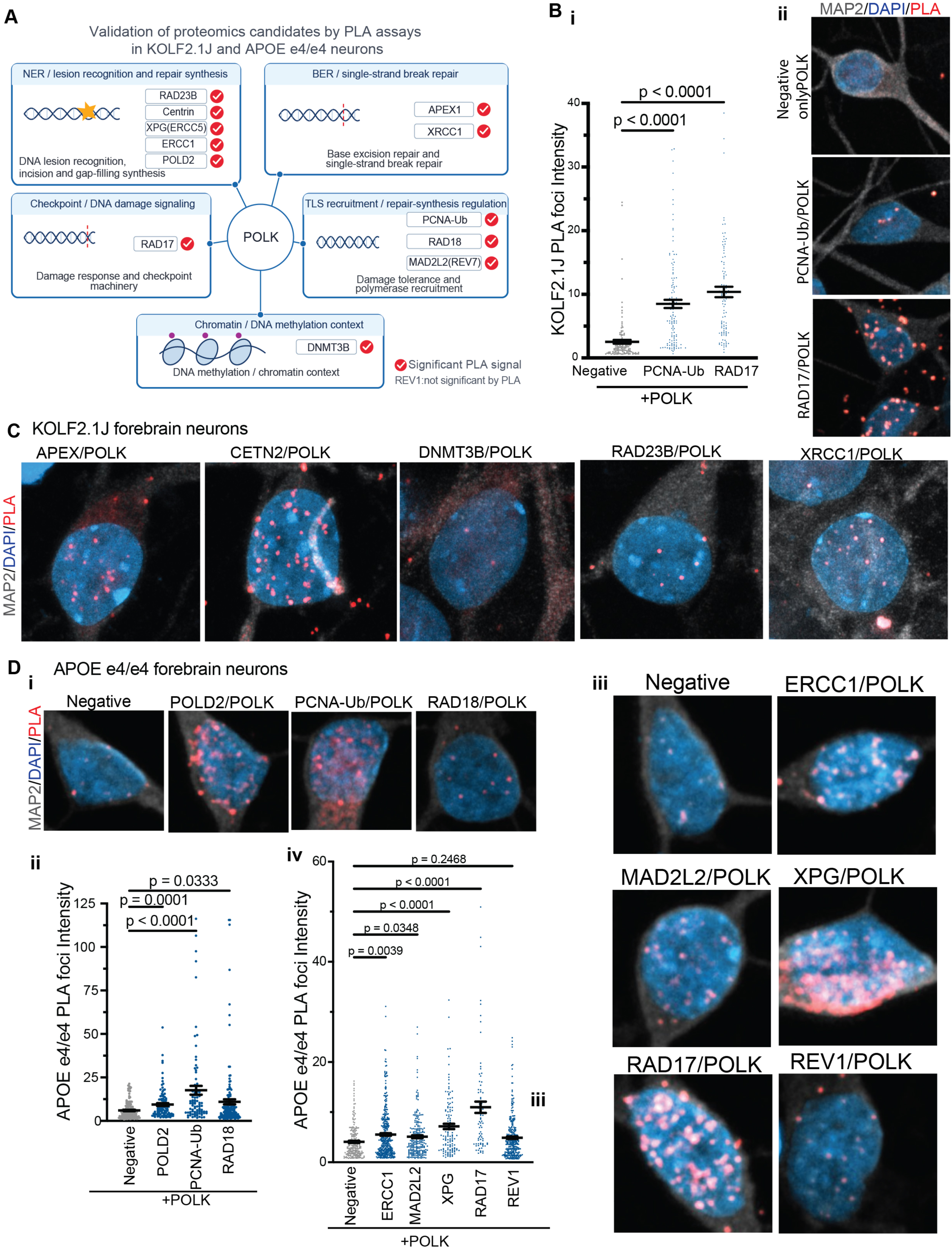
POLK is proximal to DNA repair, DNA damage-response, and chromatin-associated proteins in human forebrain neurons. (A) Exploratory nuclear POLK IP–MS candidate framework and PLA-based pathway map. Proteins shown include NER lesion recognition and repair synthesis (RAD23B, CETN2/centrin, XPG/ERCC5, ERCC1, POLD2), BER/SSBR (APEX1 and XRCC1), DNA damage response (RAD17), specialized polymerase recruitment and regulation (PCNA-Ub, RAD18, MAD2L2/REV7), and DNA methylation-associated chromatin context (DNMT3B). The nuclear IP–MS experiment had one sample per condition and was used to nominate candidates rather than to establish a statistically supported interactome. REV1 PLA was not significant. (B) KOLF2.1J forebrain neurons. (i) PLA foci intensity comparing negative control with PCNA-Ub–POLK and RAD17–POLK (P < 0.0001 for indicated comparisons). (ii) Representative MAP2/DAPI/PLA images of the negative control and PCNA-Ub–POLK and RAD17–POLK assays. (C) Representative KOLF2.1J POLK-PLA images with APEX1, CETN2, DNMT3B, RAD23B, and XRCC1; additional single-cell distributions are shown in Supplementary Fig. 3. (D) APOE e4/e4 forebrain neurons. (i) Representative negative-control, POLD2– POLK, PCNA-Ub–POLK, and RAD18–POLK PLA images. (ii) Quantification of PLA foci intensity for these candidates. (iii) Representative negative-control and ERCC1–POLK, MAD2L2–POLK, XPG– POLK, RAD17–POLK, and REV1–POLK images. (iv) Corresponding PLA foci intensity distributions; REV1 did not differ significantly from negative control (p = 0.2468).

POLK peptides were not detected by mass spectrometry. The resulting protein profile provided an initial framework for identifying candidate proteins involved in DNA repair and genome maintenance. Given the exploratory design (n = 1), we used these findings to guide the selection of biologically relevant candidates for independent validation by proximity ligation assay (PLA), rather than to establish a definitive POLK interactome.

Using KOLF2.1J and APOE e4/e4 forebrain neurons, PLA detected POLK proximity to proteins involved in distinct DNA-repair and genome-maintenance contexts (Fig. 6A–D; Supplementary Fig. 3). Within nucleotide excision repair (NER), POLK was proximal to RAD23B and centrin (CETN2), components of the XPC lesion-recognition machinery; XPG/ERCC5 and ERCC1, involved in lesion excision; and POLD2, a subunit of DNA polymerase δ involved in repair synthesis. These findings align with previous evidence that POLK and POLδ contribute to NER repair synthesis through distinct recruitment mechanisms, including ubiquitinated PCNA-dependent POLK recruitment ^12^. Notably, POLK proximity to ubiquitinated PCNA (PCNA-Ub) was detected in both KOLF2.1J and APOE e4/e4 forebrain neurons, while proximity to RAD18 further supports an association with the established RAD18-dependent PCNA ubiquitination machinery involved in POLK recruitment to damaged DNA ^17^. Beyond NER, PLA detected POLK proximity to APEX1 and XRCC1, proteins involved in base-excision and single-strand-break repair (BER/SSBR), in KOLF2.1J neurons (Fig. 6C; Supplementary Fig. S3). POLK–DNMT3B proximity was also observed (Fig. 6A,C; Supplementary Fig. S3), placing POLK near a DNA methylation-associated protein without establishing a role for DNMT3B in the repair reaction. These human-neuron proximity data extend our previous *in vivo* observations in mouse brain, in which nuclear POLK speckles were associated with BER proteins APE1 and LIG3 and with DNA-end-repair-associated proteins and damage markers ^28^. POLK was also proximal to MAD2L2/REV7, a component of polymerase ζ in translesion synthesis that also contributes to DNA-end-repair pathway choice. REV1, another TLS-associated protein, did not show a significant increase over the negative-control condition under the conditions tested (Fig. 6).

PLA detected POLK proximity to RAD17 in both KOLF2.1J and APOE e4/e4 forebrain neurons (Fig. 6B,D). RAD17 is a checkpoint clamp-loader component that recruits the RAD9–RAD1–HUS1 (9-1-1) complex and participates in ATR-dependent DNA damage signaling. RAD17 and other checkpoint-associated proteins are also expressed in postmitotic neurons, supporting functions beyond conventional cell-cycle regulation ^48^. Functional studies have further implicated RAD17 and the 9-1-1 complex in neuronal responses regulating axon regeneration ^49^. A potential connection to specialized DNA polymerases has been demonstrated in fission yeast, where recruitment of the POLK homolog DinB depends on Rad17 and the 9-1-1 complex ^50^. Although RAD17 depletion did not impair POLK recruitment during UV-induced NER in human fibroblasts ^12^, the reproducible POLK–RAD17 proximity in two human neuronal backgrounds suggests a potential checkpoint-associated context for POLK function in postmitotic neurons.

Together, the PLA measurements position neuronal POLK near proteins involved in NER lesion recognition, incision, and repair synthesis; BER/SSBR; RAD18–PCNA-Ub-mediated polymerase recruitment; checkpoint/damage-response signaling; and DNA methylation-associated chromatin environments. These findings establish spatial proximity of POLK to more than one repair complex.

### POLK depletion increases neuronal 6E10 immunoreactivity and remodels APP/endosomal, lipid, and lysosomal pathways

The exploratory proteome identified changes in APP and proteins involved in APP trafficking in POLK-knockdown KOLF2.1J neurons. APP itself showed a modest increase following POLK reduction (log2 fold change approximately +0.19) (Supplementary Table 1). We therefore asked whether POLK reduction altered APP/Aβ-associated neuronal phenotype. In KOLF2.1J-derived forebrain neurons, 6E10 immunoreactivity was low under control conditions but significantly increased following POLK depletion, with signal detected in TUJ1-positive neuronal somata and processes (Fig. 7A). APOE e4/e4 control neurons displayed substantially greater baseline 6E10 immunoreactivity than wild-type controls (Fig. 7A). Because 6E10 recognizes an N-terminal Aβ epitope that is also present in full-length APP and APP-derived fragments, these findings are interpreted as changes in 6E10 immunoreactivity rather than as direct evidence of Aβ-specific accumulation, extracellular amyloid deposition, or increased Aβ secretion.

**Figure 7.**
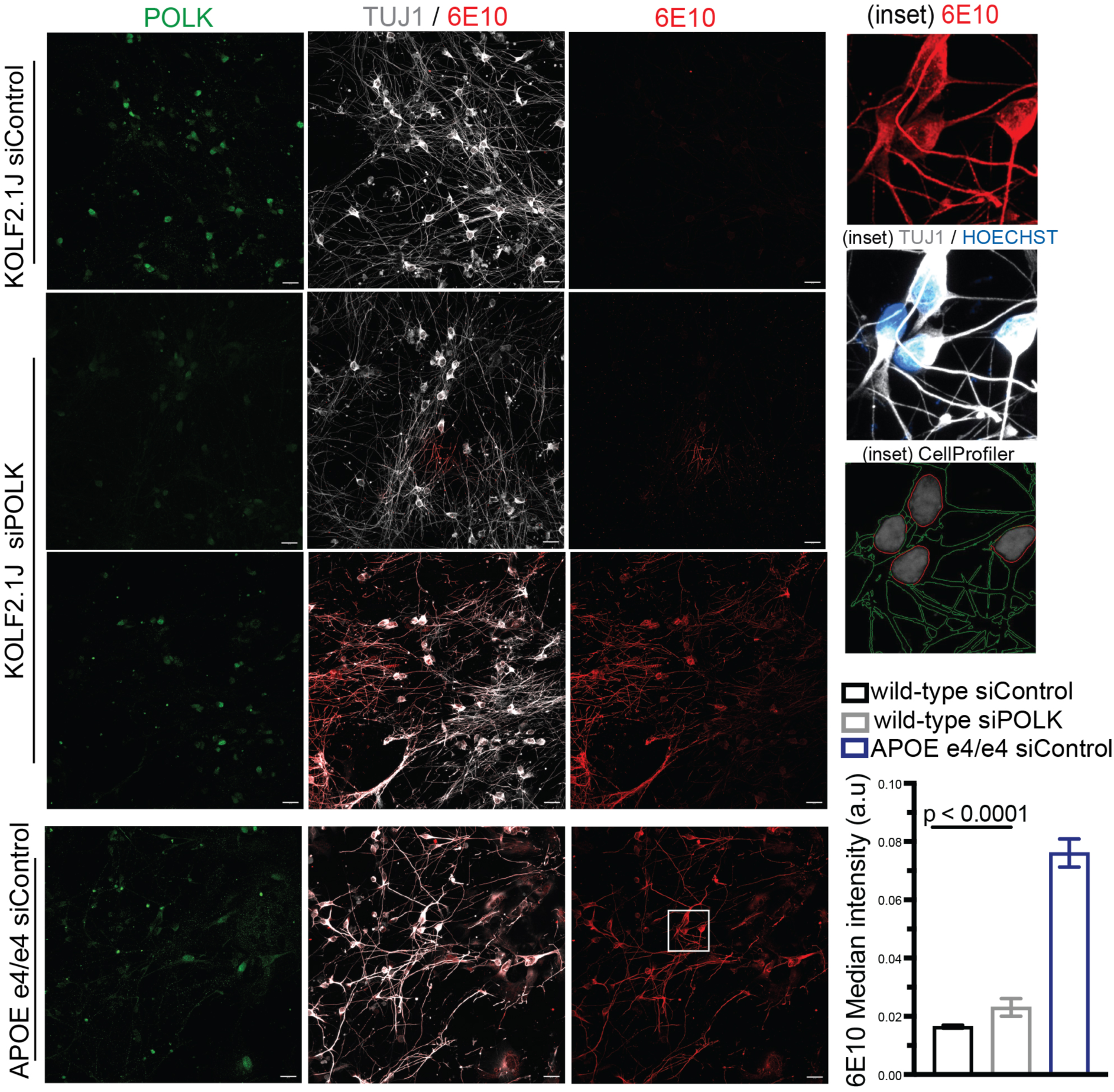
POLK depletion increases neuronal 6E10 immunoreactivity in human forebrain neurons. Representative POLK, TUJ1/6E10, and 6E10 images in KOLF2.1J siControl, KOLF2.1J siPOLK, and APOE e4/e4 siControl forebrain neurons. Magnified images show 6E10 labeling, TUJ1/Hoechst, and CellProfiler segmentation. Quantification shows significantly increased 6E10 median intensity after POLK depletion in KOLF2.1J neurons, and higher 6E10 signal in APOE e4/e4 control neurons; the indicated statistical comparison is P < 0.0001.

We next examined whether exploratory proteomics provided broader pathway context for the increased neuronal 6E10 immunoreactivity. Among 5,752 detected proteins, we selected 15 based on their established involvement in APP/Aβ biology, intracellular trafficking, lipid homeostasis, and lysosomal function. Functional annotation of this biologically selected subset highlighted APP trafficking, endocytosis, cholesterol homeostasis, and lysosomal function (Supplementary Fig. S4). Representative changes included increased RAB5B and PICALM, reduced SORL1 and NPC1, and altered abundance of lysosomal and autophagy-associated proteins. These descriptive patterns provide a pathway-level context for the increased 6E10 signal without establishing altered APP processing or pathway activity.

To determine whether these molecular changes were accompanied by AD-relevant neuronal phenotypes, we assessed tau phosphorylation and extranuclear DNA accumulation following POLK depletion in APOE e4/e4 forebrain neurons. Phosphorylation of tau at Ser396, a disease-associated epitope linked to pathological tau accumulation, was examined using a phospho-specific antibody. Somatic/cytoplasmic dsDNA was also assessed because extranuclear DNA can contribute to innate immune signaling, and our earlier proteomic analysis indicated reduced DNASE2A, a lysosomal nuclease involved in DNA degradation (Fig. 3C). POLK depletion significantly increased phospho-tau Ser396 immunoreactivity in MAP2-positive neurons (P < 0.0001; Fig. 8Ai,ii). In parallel, somatic/cytoplasmic dsDNA staining increased (P < 0.0001; Fig. 8Bi–iii), accompanied by reduced nuclear POLK (P = 0.0342; Fig. 8Biv). Together, these findings associate POLK depletion with increased disease-related tau phosphorylation and extranuclear DNA accumulation in APOE e4/e4 neurons, consistent with the altered DNA clearance machinery suggested by exploratory proteomics.

**Figure 8.**
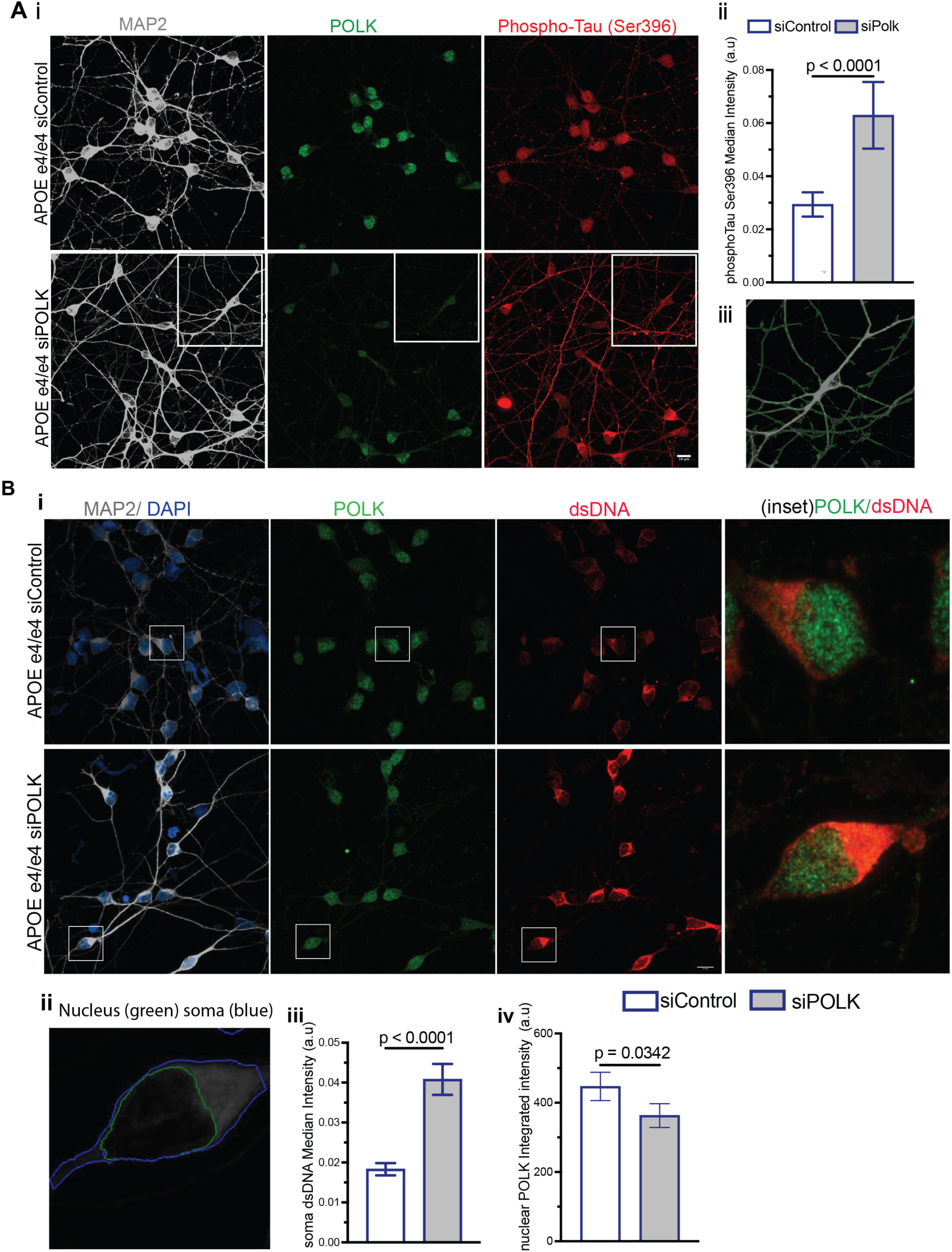
POLK depletion increases phospho-tau and cytoplasmic dsDNA in APOE e4/e4 forebrain neurons. (A) Phospho-tau Ser396 following POLK depletion in APOE e4/e4 neurons. (i) Representative MAP2, POLK, and phospho-tau Ser396 images. (ii) Quantification showing increased phospho-tau Ser396 intensity after POLK depletion (P < 0.0001). (iii) Enlarged neuronal image from the indicated region. (B) Cytoplasmic dsDNA following POLK depletion. (i) Representative MAP2/DAPI, POLK, dsDNA, and enlarged POLK/dsDNA images. (ii) Example of nucleus and soma segmentation used to quantify non-nuclear dsDNA. (iii) POLK depletion increased somatic dsDNA intensity (P < 0.0001). (iv) Nuclear POLK intensity was reduced (p = 0.0342).

### Neuronal POLK augmentation reduces p38 MAPK-associated stress

The loss-of-function experiments suggested that reduced POLK compromises neuronal genome maintenance and promotes persistent cellular stress. We therefore asked whether increasing neuronal POLK produces reciprocal effects. POLK was expressed using a neuron-directed hSyn lentiviral construct with an otherwise matched control vector (Fig. 9Ai). POLK overexpression increased neuronal and nuclear POLK immunoreactivity (p < 0.0001; Fig. 9Aii,iii).

**Figure 9.**
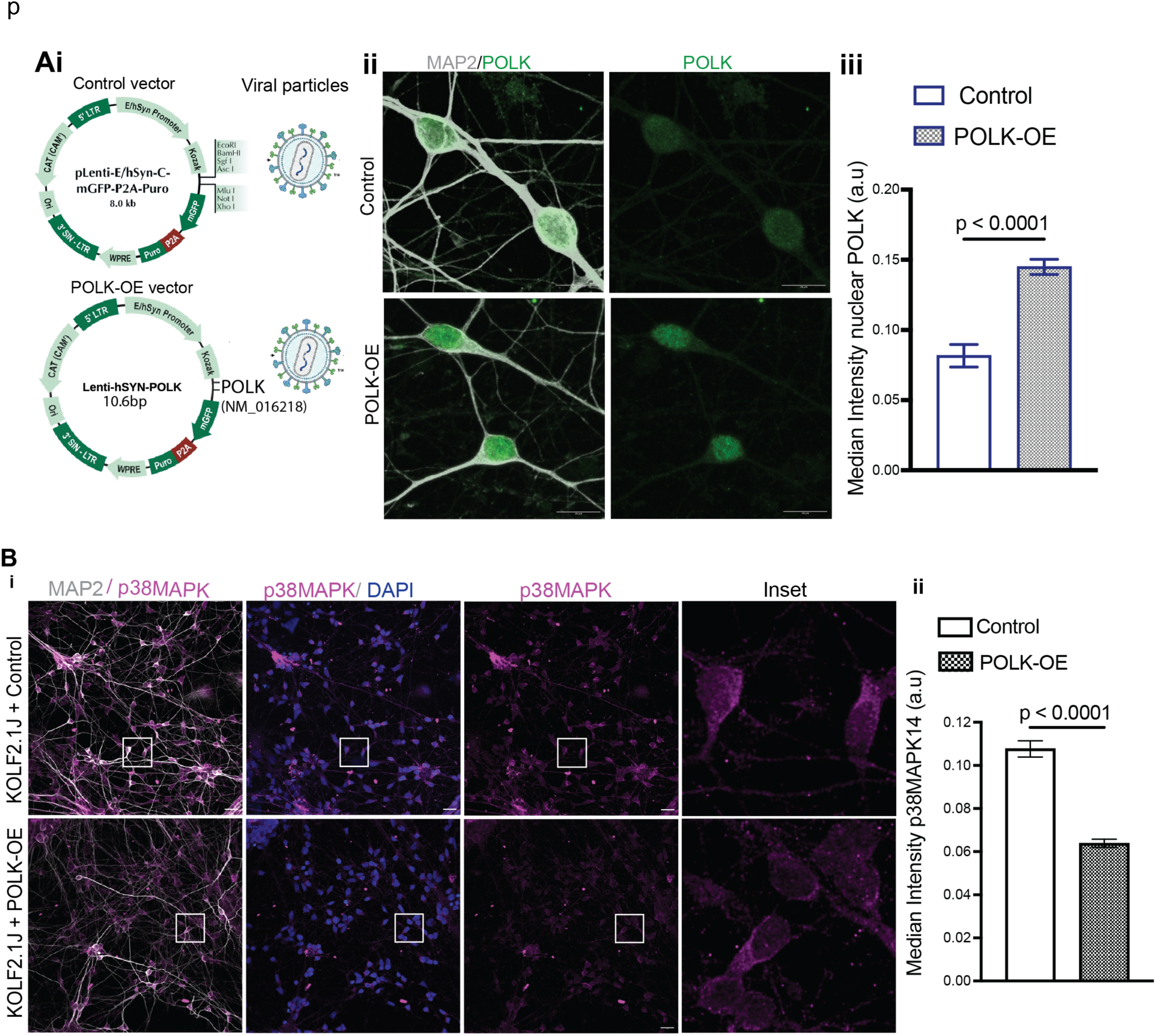
Neuronal POLK augmentation reduces p38 MAPK-associated stress. (A) Lentiviral POLK overexpression strategy. (i) Schematics of the hSyn control and hSyn-POLK lentiviral constructs and viral particles. (ii) Representative MAP2/POLK images showing increased neuronal POLK after POLK overexpression. (iii) Quantification demonstrating increased nuclear POLK (P < 0.0001). (B) p38 MAPK after POLK augmentation. (i) Representative MAP2, p38 MAPK, DAPI, and enlarged images from control and POLK-overexpressing neurons. (ii) Quantification showing reduced p38 MAPK immunoreactivity following POLK overexpression (P < 0.0001).

We first examined p38a MAPK, which increased after POLK depletion (Fig. 3F) and was nominated by MAPK14 in the exploratory proteome (Supplementary Table 1). POLK augmentation significantly reduced p38 MAPK immunoreactivity (p < 0.0001) in MAP2-positive neurons compared with control-vector cultures (Fig. 9B). Thus, p38 MAPK shows a reciprocal relationship with neuronal POLK abundance across loss-and gain-of-function conditions.

### POLK augmentation increases neuronal EdU incorporation while reducing DNA-damage-associated signals

We next examined whether POLK augmentation altered markers of DNA damage and neuronal DNA synthesis. POLK overexpression decreased 53BP1 focus intensity and reduced the proportion of neurons containing detectable 53BP1 foci from 64.1% to 46.5% (Fig. 10Ai–iii), indicating fewer neurons with focal accumulation of 53BP1, a key mediator of DNA double-strand break repair in postmitotic neurons. POLK augmentation also reduced γH2AX focus intensity (P < 0.0001; Fig. 10Bi–iii), while the proportion of neurons containing γH2AX foci remained largely unchanged (45.4% versus 44.3%). Thus, POLK overexpression reduced the prevalence and intensity of 53BP1-associated DNA repair responses while attenuating broader DNA damage-associated chromatin signaling without eliminating detectable γH2AX foci. In contrast, EdU incorporation increased following POLK augmentation (Fig. 10Ci-iii). Median EdU intensity was significantly higher (P < 0.0001), and the fraction of EdU-positive neurons increased from 16.7% to 22.7%. Together, these gain-of-function findings complement the depletion experiments, demonstrating that increased POLK abundance is associated with enhanced neuronal DNA synthesis and reduced DNA damage-associated signaling.

**Figure 10.**
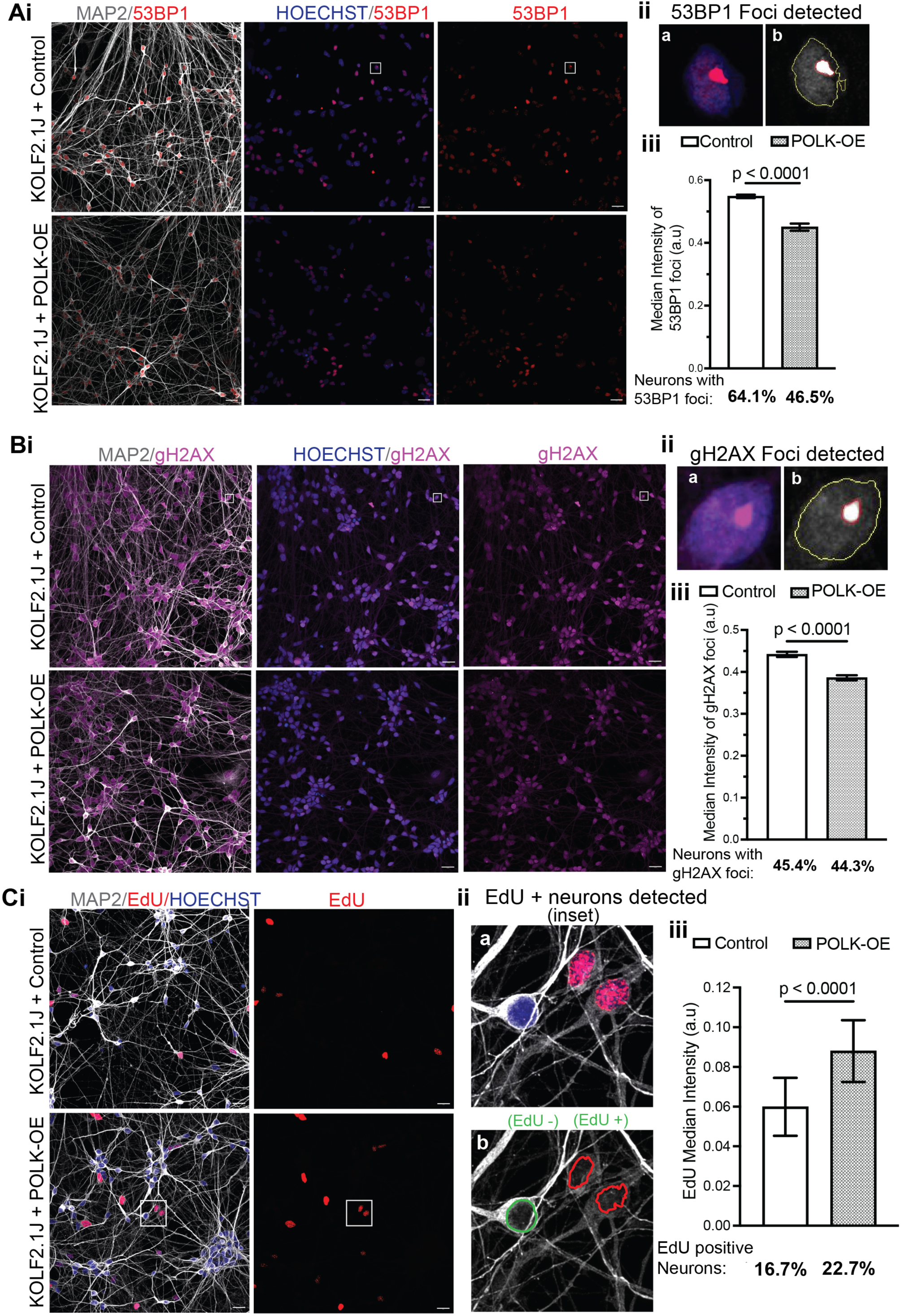
POLK augmentation increases neuronal EdU incorporation while reducing DNA-damage-associated signals. (A) 53BP1 analysis after POLK overexpression. (i) Representative MAP2/53BP1, Hoechst/53BP1, and 53BP1 images. (ii) Example of automated 53BP1-focus detection. (iii) POLK overexpression reduced 53BP1 focus intensity (P < 0.0001); the proportion of neurons with detectable 53BP1 foci decreased from 64.1% to 46.5%. (B) γH2AX analysis. (i) Representative MAP2/γH2AX, Hoechst/γH2AX, and γH2AX images. (ii) Example of automated γH2AX-focus detection. (iii) γH2AX focus intensity was reduced by POLK overexpression (P < 0.0001); the fraction of neurons with detectable γH2AX foci was 45.4% in control and 44.3% after POLK overexpression. (C) EdU incorporation. (i) Representative MAP2/EdU/Hoechst and EdU images. (ii) Examples of EdU-positive and EdU-negative neuronal segmentation. (iii) POLK overexpression increased EdU median intensity (P < 0.0001), with EdU-positive neurons increasing from 16.7% to 22.7%.

### POLK augmentation reduces cytoplasmic dsDNA and AD-relevant phenotypes in APOE e4/e4 neurons

Finally, we asked whether increasing POLK could modify the AD-relevant phenotypes observed in APOE e4/e4 neurons. POLK overexpression significantly reduced somatic dsDNA intensity (P < 0.0001; Fig. 11Ai,ii), providing a reciprocal counterpart to the increase in cytoplasmic dsDNA after POLK depletion (Fig. 8B). POLK augmentation also produced a modest but significant reduction in 6E10 immunoreactivity (p = 0.0133; Fig. 11Bi,ii) and a larger reduction in phospho-tau Ser396 (P < 0.0001; Fig. 11Ci,ii). Representative imaging with the AT8 antibody recognizing phospho-tau Ser202/Thr205 similarly showed lower phospho-tau signal in POLK-overexpressing neurons (Fig. 11Di,ii), although this panel provides qualitative support.

**Figure 11.**
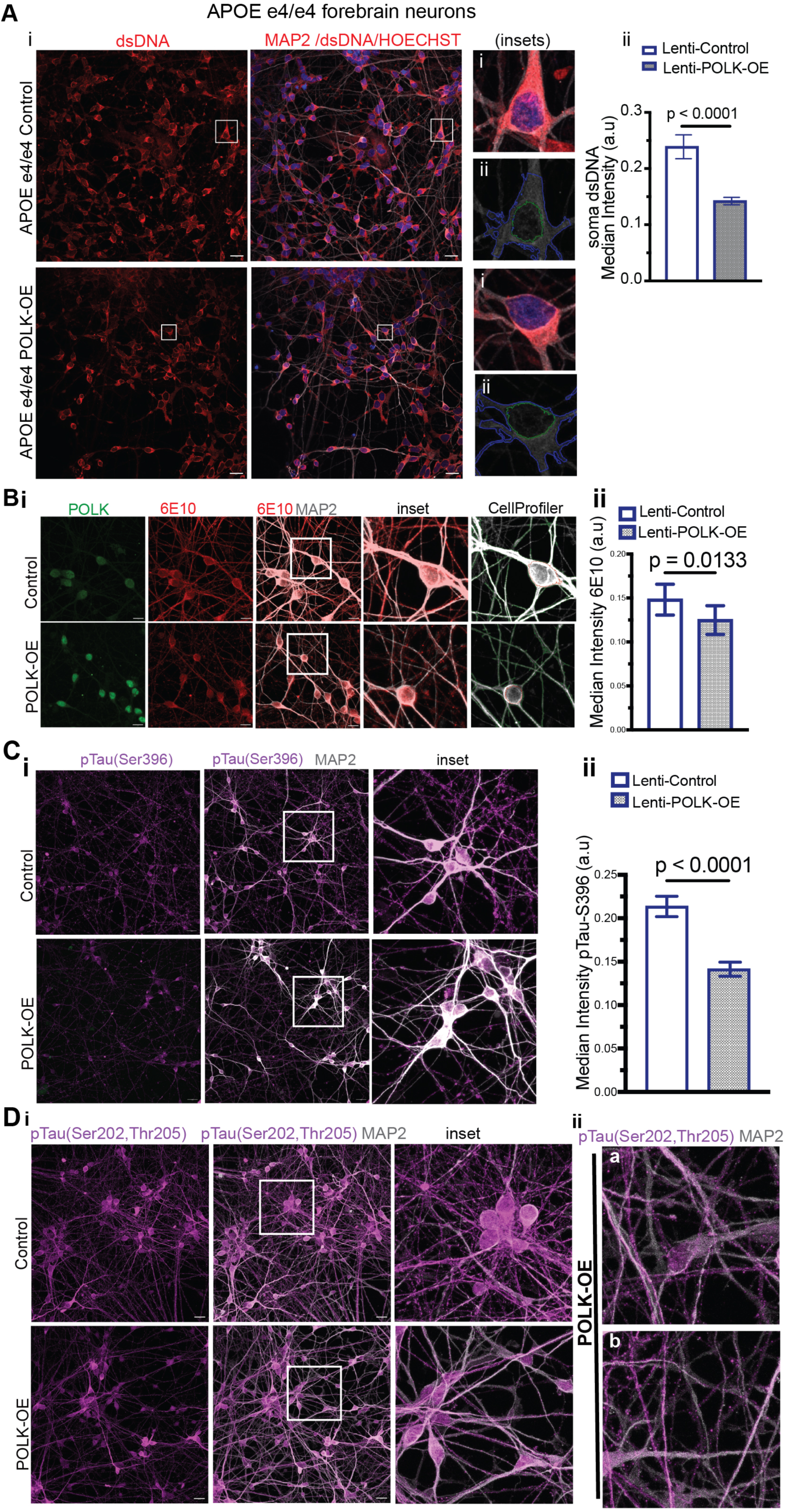
POLK augmentation reduces cytoplasmic dsDNA and AD-relevant phenotypes in APOE e4/e4 neurons. (A) Somatic dsDNA in APOE e4/e4 forebrain neurons after POLK overexpression. (i) Representative dsDNA and MAP2/dsDNA/Hoechst images with enlarged examples and soma/nuclear segmentation. (ii) POLK augmentation significantly reduced somatic dsDNA intensity (P < 0.0001). (B) 6E10 immunoreactivity. (i) Representative POLK, 6E10, MAP2, enlarged, and CellProfiler-segmented images in control and POLK-overexpressing APOE e4/e4 neurons. (ii) Quantification showing reduced 6E10 intensity following POLK overexpression (p = 0.0133). (C) Phospho-tau Ser396. (i) Representative phospho-tau Ser396 and MAP2 images. (ii) POLK overexpression significantly reduced phospho-tau Ser396 intensity (P < 0.0001). (D) Phospho-tau Ser202/Thr205 (AT8) staining. (i) Representative control and POLK-overexpressing cultures with enlarged regions. (ii) Higher-magnification POLK-overexpressing neurons. AT8 images provide qualitative evidence of reduced phospho-tau signal.

Together, the loss- and gain-of-function experiments establish reciprocal relationships between neuronal POLK abundance, DNA synthesis, and genome-maintenance-associated phenotypes. POLK depletion reduces DNA synthesis in primary mouse cortical neurons and human forebrain neurons, increases DNA damage and stress-associated signaling, alters nuclear and organelle homeostasis, and promotes AD-relevant cellular changes. Conversely, POLK overexpression increases neuronal DNA synthesis while reducing 53BP1 and γH2AX signals, p38α MAPK immunoreactivity, cytoplasmic dsDNA accumulation, 6E10 immunoreactivity, and tau phosphorylation. These complementary findings support a model in which POLK contributes to genome maintenance in postmitotic CNS neurons through repair-associated DNA synthesis, highlighting its relevance to neuronal resilience during aging and in AD-risk contexts.

## DISCUSSION

Our findings identify POLK as a regulator of ongoing DNA synthesis and genome-maintenance-associated responses in postmitotic CNS neurons, extending the functional relevance of a specialized TLS polymerase beyond its established roles in proliferating cells. Endogenous POLK localized near newly synthesized DNA, POLK depletion reduced EdU incorporation in primary mouse cortical neurons and two independently derived human forebrain neuronal lines, and POLK augmentation increased EdU incorporation in human neurons. Importantly, these relationships were observed during ongoing neuronal DNA synthesis without an experimentally imposed genotoxic challenge. Changes in POLK abundance were also accompanied by reciprocal DNA damage-associated responses, together with nuclear and organelle remodeling, senescence-like stress, and AD-relevant cellular phenotypes. The convergence of localization, loss-of-function, and gain-of-function findings identifies POLK as an important candidate component of the repair-synthesis machinery supporting genome maintenance in postmitotic CNS neurons and connects this fundamental process to neuronal vulnerability (Fig. 12).

**Figure 12.**
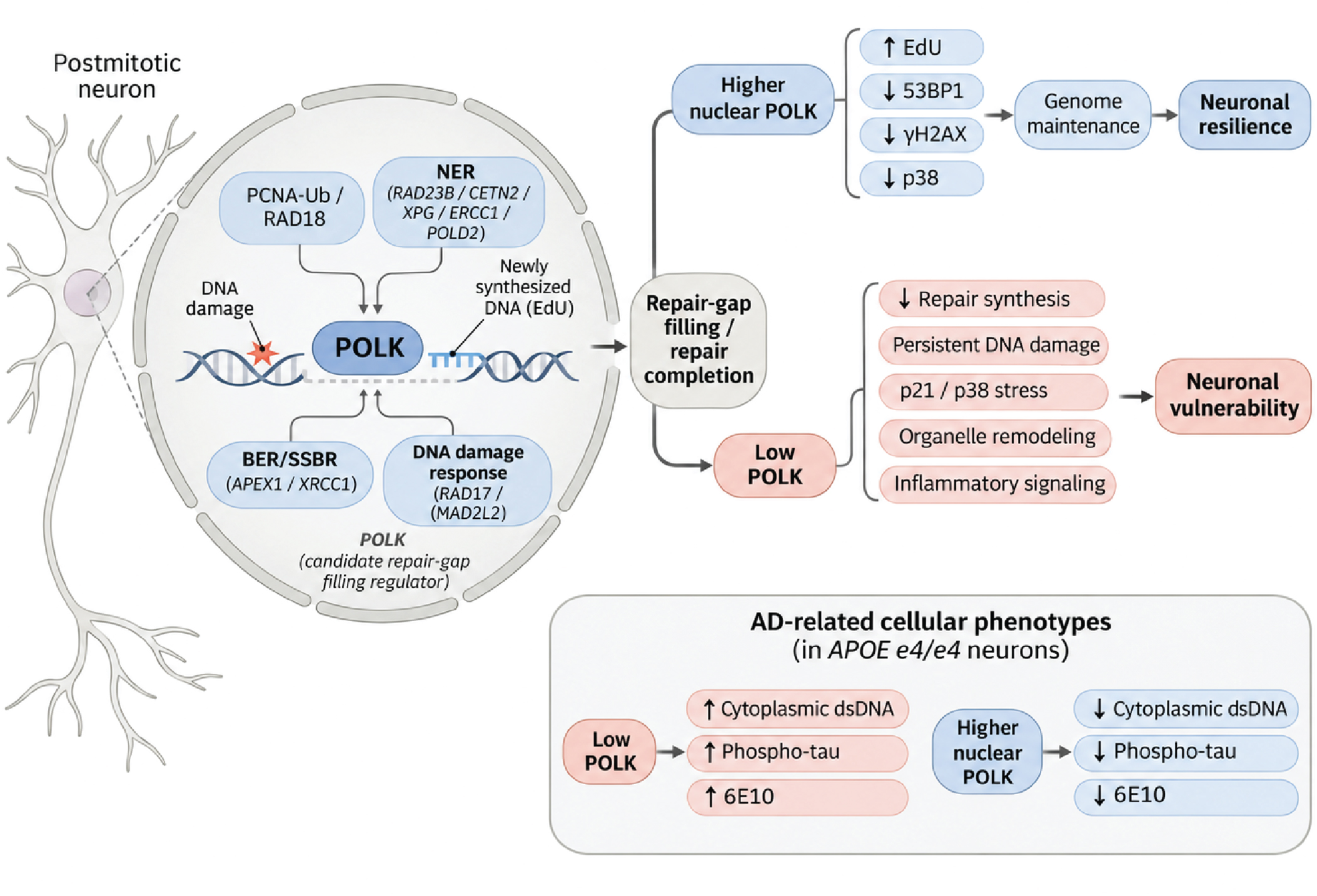
Poposed Working Model

Genome-wide Repair-seq and SAR-seq studies established recurrent repair-associated DNA synthesis in postmitotic human neurons, particularly at active genomic regions and enhancers ^6,47^. These studies mapped neuronal repair synthesis and identified contributions from established repair mechanisms, while the involvement of specialized TLS polymerases in ongoing repair synthesis in postmitotic CNS neurons remained unresolved. POLK was a particularly relevant candidate because of its established contribution to NER pathway DNA synthesis in human cells and cisplatin-induced repair synthesis in postmitotic dorsal root ganglion neurons ^12,27^. The proximity of endogenous POLK to EdU-labeled nascent DNA, reduced EdU incorporation following POLK depletion in mouse cortical and independently derived human forebrain neurons, and increased incorporation following POLK augmentation now extend this connection to ongoing DNA synthesis in postmitotic CNS neurons. The SIRF and imaging measurements position POLK near nascent DNA, while reciprocal perturbations establish a relationship between POLK abundance and neuronal nucleotide incorporation. These observations expand the candidate polymerase repertoire for neuronal gap filling beyond POLβ-dependent short-patch repair, whose impairment shifts synthesis toward longer repair tracts ^6^. Genome-wide mapping of POLK-sensitive synthesis will help define whether specialized polymerases support particular neuronal loci, including enhancers and transcriptionally active regions with persistent repair demand.

NER provides a compelling mechanistic context for these findings. In human forebrain neurons, POLK was proximal to factors operating at successive stages of NER, including lesion recognition (RAD23B and centrin/CETN2), incision (XPG/ERCC5 and ERCC1), and repair synthesis (POLD2). POLK was also proximal to ubiquitinated PCNA (PCNA-Ub) in KOLF2.1J and APOE e4/e4 neurons, together with RAD18, which promotes PCNA ubiquitination and POLK recruitment at damaged DNA in proliferating cells ^12,17^. The convergence of recognition, incision, synthesis, and recruitment-associated factors in the neuronal POLK proximity landscape provides a coherent molecular framework for specialized gap filling in postmitotic neurons. Recent evidence that incomplete NER gap filling can sustain neuronal strand breaks underscores the importance of completing this synthesis step ^10^. These findings raise the possibility that RAD18-dependent PCNA ubiquitination regulates POLK recruitment to sites of ongoing neuronal repair synthesis. Examining this mechanism through RAD18–PCNA perturbation, POLK–nascent DNA proximity, and repair-synthesis measurements could establish how specialized polymerase recruitment contributes to genome maintenance in postmitotic neurons.

The increase in neuronal p21 following POLK depletion adds a potential regulatory dimension to this recruitment mechanism. In proliferating human cells, p21 modulates access of specialized polymerases, including POLK and PrimPol, to PCNA during nascent DNA synthesis, with consequences for genome stability ^51^. In our neurons, increased p21 coincided with reduced EdU incorporation, whereas proximity between POLK and PCNA-Ub was independently detected. These observations suggest a testable feedback relationship in which a POLK-sensitive stress response could influence access of specialized polymerases to neuronal repair intermediates. Determining whether p21 alters POLK recruitment or the efficiency of repair synthesis would link a prominent senescence-like response in our models to a defined genome-maintenance mechanism.

Our findings also place POLK in molecular environments extending beyond NER. POLK has established functions in strand-break repair, replication-independent interstrand crosslink repair, and synthesis across structurally challenging DNA templates ^13–15^. In our earlier aging-mouse study, nuclear POLK was associated with BER/SSBR and DNA-end-repair proteins and DNA damage markers ^28^. Here, proximity to APEX1 and XRCC1 associates neuronal POLK with base-excision and single-strand-break repair machinery, while MAD2L2/REV7 links it to polymerase ζ-associated damage tolerance and DNA-end-repair pathway choice. REV1 did not show significant PLA enrichment over the negative control under the conditions tested, indicating that elevated POLK proximity is not a uniform feature of all TLS-associated factors. Together, these observations support investigating whether POLK participates in distinct repair assemblies according to lesion type, gap structure, or neuronal chromatin context, rather than operating exclusively through one canonical TLS complex.

POLK–DNMT3B proximity opens an additional direction linking specialized DNA synthesis to methylation-associated chromatin maintenance. Repair-seq and SAR-seq identified neuronal repair synthesis at defined genomic regions, including enhancers and CpG-associated sites linked to epigenetic regulation ^6,7,47^. Subsequent work established TET–TDG-mediated active DNA demethylation as a source of neuronal single-strand breaks repaired through both short-and long-patch synthesis ^52^. DNMT3B functions in DNA methylation rather than demethylation, making its proximity to POLK particularly interesting in the context of repair-associated chromatin remodeling. Together with POLK proximity to APEX1 and XRCC1, this finding raises the hypothesis that specialized POLK-associated synthesis may occur at demethylation-linked repair intermediates and coordinate with subsequent maintenance or remodeling of local methylation states. Joint mapping of POLK-dependent repair synthesis, DNA methylation, and regulatory-region activity would directly test this possibility and connect neuronal genome repair to epigenetic stability.

Additional proximity findings broaden the genomic settings in which neuronal POLK may act. POLK–RAD17 proximity in both KOLF2.1J and APOE e4/e4 forebrain neurons places POLK near checkpoint-associated DNA damage-response machinery, suggesting that surveillance of DNA repair intermediates may be coordinated with specialized polymerase recruitment even in postmitotic cells. This provides a tractable hypothesis for testing whether RAD17-associated signaling influences POLK localization and synthesis at persistent repair gaps. Transcription-associated R-loops represent a complementary direction, particularly given recent evidence that POLK recruits DDX23 to promote R-loop resolution ^53^. In neurons, where transcription and topoisomerase activity generate recurring DNA lesions ^2,3,8^, it will be important to determine whether POLK-dependent DNA synthesis is enriched at R-loop-associated or other transcriptionally stressed loci. These hypotheses connect the observed neuronal proximity landscape to defined pathways that can be separated experimentally by recruitment perturbations and genomic localization of repair synthesis.

The cellular consequences of POLK depletion underscore the broader importance of maintaining adequate neuronal DNA-synthesis capacity. Increased nuclear area and lamin A/C remodeling occurred alongside elevated senescence-associated β-galactosidase activity, altered lysosomal staining, reduced MitoTracker signal, and increased p21, inflammatory transcripts, and nuclear p38 MAPK immunoreactivity (Figs. 1–3). These coordinated changes resemble multidimensional senescence-like stress responses described in postmitotic neurons following persistent DNA damage ^42,44^. The reduction in p38 MAPK immunoreactivity following POLK augmentation (Fig. 9) provides an important reciprocal response. Together, the phenotypes indicate that neuronal POLK abundance is linked not only to DNA synthesis but also to nuclear organization, organelle homeostasis, and stress signaling. They lead to the testable hypothesis that diminished capacity to complete repair-associated synthesis contributes to the persistence of damage-associated responses, which may in turn reinforce inflammatory and senescence-like neuronal states. Measuring repair completion after POLK depletion and catalytic rescue will be important for establishing the sequence connecting DNA synthesis to these broader phenotypes.

The complementary responses in APOE e4/e4 forebrain neurons extend the significance of these findings to an AD-risk genetic context. POLK depletion increased phospho-tau Ser396 and somatic/cytoplasmic double-stranded DNA, while POLK augmentation reduced both readouts and lowered 6E10 immunoreactivity (Figs. 7, 8, and 11). Increased neuronal EdU incorporation and reduced 53BP1 and γH2AX focus intensity following POLK augmentation (Fig. 10) connect these disease-relevant readouts to a favorable shift in genome-maintenance-associated responses. Because 6E10 recognizes epitopes shared by amyloid-β and APP-derived species, the observed changes reflect APP/Aβ-related immunoreactivity rather than direct measurement of amyloid production or deposition. The exploratory reduction in DNASE2A also suggests a potential intersection between genome maintenance and cytoplasmic DNA clearance. Collectively, these findings support testing whether improved repair synthesis can reduce persistent damage-associated signaling and downstream tau, APP-related, or cytoplasmic DNA phenotypes. Such experiments could establish whether neuronal genome maintenance represents a mechanistic point of convergence between aging-associated stress and AD-relevant cellular vulnerability.

### Limitations

Our results connect POLK abundance to ongoing neuronal DNA synthesis and multiple genome-maintenance-associated phenotypes; the specific POLK-catalyzed repair reactions and genomic substrates remain to be defined. EdU, SIRF, and PLA measure incorporation or proximity rather than direct enzymatic gap filling or protein binding. The unreplicated proteomic and IP–MS datasets were used to nominate pathways, with selected associations independently evaluated by PLA. hiPSC-derived neurons model postmitotic neuronal and AD-risk genetic contexts without reproducing all aspects of aging or disease. Catalytic and recruitment-defective rescue, repair-kinetic measurements, and genome-wide synthesis mapping provide direct next steps for establishing mechanism and biological consequence.

Together, our findings identify POLK as a regulator of ongoing DNA synthesis and genome-maintenance-associated responses in postmitotic CNS neurons and establish a foundation for understanding how specialized TLS polymerases contribute to repair outside DNA replication. The reciprocal depletion and augmentation phenotypes, together with proximity to NER, BER/SSBR, PCNA-Ub, checkpoint, and chromatin-associated factors, point to several experimentally tractable routes through which POLK may help maintain neuronal genome integrity. Considered alongside our earlier observation of age-associated loss of nuclear POLK ^28^, these results support the hypothesis that reduced availability of specialized repair-synthesis machinery contributes to persistent neuronal stress and disease-relevant vulnerability (Fig. 12). Testing where and how POLK is recruited, and whether restoration of its repair-associated activity improves neuronal responses to aging-and AD-related stress, will connect this molecular mechanism to a broader understanding of neuronal genome maintenance and neurodegenerative disease.

## METHODS

### Primary Cortical Neuronal Culture, siRNA Transfection, and Lentiviral Transduction

Primary cortical neuronal cultures were prepared from postnatal day 0–1 (P0–P1) mice using the Neural Tissue Dissociation Kit – Postnatal Neurons (Miltenyi Biotec, Cat. No. 130-094-802), according to the manufacturer’s instructions, with minor modifications. Briefly, 24-well culture plates and 8-well chamber slides (Nunc Lab-Tek II CC2 Chamber Slide System, Thermo Scientific, Cat. No. 154941) were coated with 50 µg/mL poly-D-lysine (Thermo Scientific, Cat. No. A3890401) for 1h at room temperature. The coated surfaces were washed three times with sterile distilled water and allowed to dry under sterile conditions.

Following humane euthanasia, brains were collected from P0–P1 mouse pups in ice-cold Dulbecco’s phosphate-buffered saline supplemented with calcium and magnesium (DPBS+). The cerebral cortices were carefully dissected under a stereomicroscope and collected for tissue dissociation. Cortical tissues were enzymatically dissociated at 37°C using the Neural Tissue Dissociation Kit – Postnatal Neurons, according to the manufacturer’s protocol. The resulting tissue suspension was gently triturated using a 1-mL pipette tip to obtain a single-cell suspension and passed through a 70-µm cell strainer. The filtrate was centrifuged at 300 × g for 10 min, and the resulting cell pellet was resuspended in complete neuronal culture medium.

The cell suspension was quantified, and cells were seeded onto poly-D-lysine-coated 24-well plates and 8-well chamber slides in complete neuronal culture medium consisting of Neurobasal™ Medium supplemented with 2% B-27™ Supplement, 0.5 mM GlutaMAX™, and 1% penicillin–streptomycin. Cultures were maintained at 37°C in a humidified incubator containing 5% CO₂. Twenty-four hours after plating, cytosine arabinoside (Ara-C; 2.5 µM; Sigma-Aldrich, Cat. No. C1768) was added to the cultures to inhibit glial cell proliferation. Half of the culture medium was replaced with fresh complete neuronal culture medium every 2–3 days.

On day in vitro 10 (DIV10), primary cortical neuronal cultures were subjected to siRNA transfection and lentiviral shRNA transduction, as previously described ^28^.

### hiPSC to forebrain differentiation

Matrigel (Corning 354277) was thawed on ice overnight, and using cooled P200 tips, Matrigel was diluted in DMEM/F12 with 15 mM HEPES (Catalog 36254) and applied to six-well plates. Two iPSC lines KOLF2.1J (Cat#JISC1000) as wild type and APOE C112R (E4/E4) [single nucleotide variant (SNV)/SNV, Cat#JISC1150] obtained from The Jackson Laboratory were seeded as single cells on six-well plates in mTeSR™ Plus (Catalog 100-0276, Stemcell Technologies) medium supplemented with 10 µM ROCK inhibitor Y-27632 and cells were allowed to adhere for 24 h at 37°C and 5% CO₂, medium was replaced after 24 h. ReLeSR™ (#100-0484) was used for further passaging iPSCs. iPSCs were maintained for several days and subsequently passaged into single cells and cultured in STEMdiff™ SMADi Neural induction medium for 21 days; cells were dissociated to single cells with Accutase™ on days 6, 12, and 18. On day 19, the medium was changed to STEMdiff™ Forebrain Neuron Differentiation medium and through day 24, the medium was changed daily with the Forebrain Neuron Differentiation medium. Cells were subsequently passaged and maintained in STEMdiff™ Forebrain Neuron Maturation medium for a few days, and full medium was changed every other day.

The KOLF2.1J parental hiPSC line is designated JIPSC1000_KOLF2.1J (The Jackson Laboratory, strain no. 765000); the engineered homozygous APOE e4/e4 line is JIPSC1150_APOE_C112R_SNV/SNV (strain no. 765075). These lines were differentiated through a neural progenitor stage into forebrain neurons using STEMCELL Technologies reagents and the manufacturer’s protocols.

### NPC to forebrain differentiation

Human iPSC-Derived Neural Progenitor Cells (200-0620) were grown according to the manufacturer’s instructions with some modifications. NPCs were expanded with Neural Progenitor Medium (05833) for several days and passaged with AccutaseTM (07920), cells were switched to Forebrain Neuron Differentiation (#08600) for 7 days and cell were further passaged onto six-well plates coated with polyornithine (# P4957) and Laminin (#L2020) and the medium was changed to BrainPhys with supplements (#08605) for 14 days. Cells were further used for experiments.

The neural progenitor cells were purchased directly from STEMCELL Technologies (catalog no. 200-0620) and were derived from the SCTi003-A human iPSC line (RRID:CVCL_C1W7), originally generated from peripheral blood mononuclear cells of a healthy female donor. These commercially sourced CNS-type NPCs were differentiated into SCTi003-A-derived forebrain neurons according to the manufacturer’s protocol, using the STEMdiff Forebrain Neuron Differentiation Kit (catalog no. 08600) and the STEMdiff Forebrain Neuron Maturation Kit (catalog no. 08605). The resulting neuronal cultures were assayed at differentiation days 55 and 73, as shown in Fig. 5.

### SiRNA

Neurons underwent siRNA transfection with Dharmacon Accell technology. Accell SMARTpool siRNA POLK was diluted to a stock concentration of 100 μM in 1X siRNA Buffer (Dharmacon horizon discovery # E-021038-00) and then diluted to 1 μM in Accell siRNA delivery media. An Accell non-targeting siRNA pool was used as a negative control (# D-001910-10). Between days 45 and 50, neuron maintenance media was removed and the Accell siRNA delivery media was added for 72 h. After 72 h, cells were fixed with 4% paraformaldehyde, and immunofluorescence assays were performed.

The POLK Accell SMARTpool comprised four POLK-targeting siRNAs; the non-targeting Accell pool served as the matched siRNA control. Matched siControl and siPOLK conditions were applied to multiple separately treated culture wells or chamber-slide chambers in the imaging assays. Where indicated, independent treatments and neuronal culture preparations were analyzed across experiments.

### Lentivirus for Overexpression

POLK expression plasmid RC219806L4 and the OriGene PS100115 lentiviral gene-expression vector were first transformed into DH5α competent E. coli. Following transformation, the bacteria were plated on LB agar containing chloramphenicol for selection, and individual colonies were subsequently expanded in LB broth containing chloramphenicol. Plasmid DNA from the bacterial cultures was purified using the Qiagen Plasmid Maxi Kit. The purified RC219806L4 and PS100115 plasmids were then subjected to restriction digestion using AsiSI and MluI restriction enzymes to prepare the POLK insert and linearized PS100115 vector for cloning. The desired DNA fragments were separated by agarose gel electrophoresis and purified using the Monarch DNA Gel Extraction Kit. The POLK insert was subsequently ligated into the prepared PS100115 lentiviral vector using the Quick Ligation Kit. Following ligation, the recombinant plasmid was transformed into competent E. coli, and individual colonies were selected and expanded for plasmid DNA preparation. The resulting plasmid DNA was used for lentiviral vector production. The POLK-containing lentiviral construct was introduced into HEK293T cells by transfection, following the OriGene protocol with modifications, and the resulting lentivirus was subsequently collected for use in POLK overexpression experiments.

The human POLK open reading frame corresponds to transcript NM_016218. The neuron-directed expression construct was designated Lenti-SYN-POLK (POLK-OE), using the OriGene pLenti-E/hSyn-C-mGFP-P2A-Puro vector (PS100115) with the human synapsin I promoter. The parental hSyn-mGFP-P2A-Puro plasmid was used as the matched GFP-expressing control. Correct POLK insert integration and orientation were assessed by restriction digestion and agarose gel electrophoresis. The POLK-overexpression and control vectors were prepared and compared in parallel.

Replication-incompetent viral particles were produced using the OriGene Lentiviral Packaging Kit (SKU TR30037) in the recommended packaging cell system, following the manufacturer’s instructions. Viral supernatant was harvested, cleared of cellular debris by centrifugation, and concentrated with Lenti-X Concentrator (Takara Bio, catalog no. 631231). Concentrated virus was aliquoted and stored at −80 °C until use. The working viral input was established empirically in human forebrain neurons by evaluating GFP or POLK expression together with neuronal survival and the preservation of MAP2-or TUJ1-positive morphology.

### Transduction

For viral transduction, neurons were plated on Lab-Tek II chamber slides and POLK-overexpression virus was added according to the experimentally optimized multiplicity of infection (MOI) to the medium for 20 h; cultures were maintained at 37°C and 5% CO₂, after which the medium was replaced with conditioned medium and cells were maintained for 5 days and media was changed with fresh media once after 3 days and then fixed with 4% PFA for immunofluorescence staining as described above.

For POLK augmentation experiments, mature KOLF2.1J-and APOE e4/e4-derived forebrain neurons were transduced with the Lenti-SYN-POLK virus or the parental GFP-expressing control virus under matched culture conditions. Transduced cultures were assessed by POLK immunofluorescence and neuronal markers after the expression period.

### Immunofluorescence of primary cortical neurons

At 48 h after siRNA treatment and 72 h after shRNA transduction, primary cortical neurons were washed with DPBS+ and fixed with 4% paraformaldehyde (PFA) for 20 min at room temperature. Cells were subsequently permeabilized with 0.3% Triton X-100 for 5 min. After permeabilization, cells were blocked for 1 h in DPBS+ containing 5% goat serum and 0.3% Triton X-100. Cells were then incubated overnight at 4°C with the appropriate primary antibodies (Table). The following day, cells were washed and incubated with the corresponding fluorescently labelled secondary antibodies (Table) for 1 h. Nuclei were counterstained with Hoechst 33342 (Thermo Fisher Scientific). Samples were mounted using mounting medium, stored at 4°C, and imaged using a Zeiss LSM confocal microscope. Image analysis was performed using CellProfiler ^54^(version 4.2.6) and ImageJ (NIH). Quantitative analyses were performed using at least three independent biological replicates.

### Immunofluorescence assays

Acidic lysosomes were stained with LysoTracker™ Red DND-99 (Thermo Fisher Scientific, USA; Cat. No. L12941). MitoTracker® Red CMXRos (Invitrogen, M7512) was used for mitochondrial staining according to the manufacturer’s instructions. Neuron cultures were washed three times with DPBS and were fixed using 4% w/v paraformaldehyde for 20 min and washed three times in PBS for 5 min. Blocking was done with 10% goat serum and 0.5% Triton X-100 for 1 h. Following this blocking step, samples were incubated overnight at 4°C with primary antibodies. Primary antibodies included mouse anti Polk (1:200, sc-166667), mouse anti beta III Tubulin antibody [2G10] (1:500, AB7808-1001), mouse anti-β-Amyloid (1:500, SIG-39320), rabbit Phospho-Tau (Ser396) (1:500, 44-752G), mouse Rev I (1:200, SC-393022), rabbit CETN2 centrin 2 (1:300, 15877-1-AP), rabbit RAD23B (1:300, 12121-1-AP), rabbit XRCC6 (1:300, A7330), rabbit APEX1 (1:300, A11238), rabbit gH2AX (1:1000, NB 100-384), rabbit 53BP1 (1:300, NB100-304), chicken MAP2 (1:1000, ab92434), rabbit APP Antibody (6E10) (1:500, NBP2-62566), rabbit APOE (1:200, 18254-1-AP), rabbit XPG (1:100, 11331-1-AP), rabbit MAD2L2 (1:100, 12683-1-AP), rabbit RAD17 (1:100, 13358-1-AP), rabbit ERCC1 (1:50, 13358-1-AP), rabbit POLD2 (1:200, 29765-1-AP), chicken GFAP (1:500, 4937975), rabbit Ubiquityl-PCNA (1:200, #13439), rabbit RAD18 (1:100, 183331-1-AP). Following incubation with corresponding secondary antibodies Alexa fluor (1:1000 dilution Invitrogen) for 1 h, the samples were washed three times with DPBS. Finally, samples were incubated for 5 min with Hoechst 33342 solution (1:1,000 dilution in DI H2O, Invitrogen), followed by mounting with Fluromount-GTM and coverslips (#2975-246) (Details of Antibodies in supplementary sheet). Slides were imaged using a Zeiss LSM 900 confocal microscope

Additional antibodies and imaging reagents used in the POLK-augmentation experiments included phospho-tau Ser202/Thr205 (AT8; Addgene, catalog no. 216150), double-stranded DNA (clone AE-2; Sigma-Aldrich, MAB1293, 1:300), and p38 MAPK (Proteintech, 66234-1-Ig, 1:100). The dsDNA assay was evaluated in neuronal somata after excluding the Hoechst-defined nuclear compartment. Anti-6E10 immunoreactivity was interpreted as detection of APP/Aβ-related epitopes, not as a specific measurement of amyloid-β production or extracellular deposition.

### Microscopy acquisition and quantitative image analysis

Confocal imaging was performed using a Zeiss LSM 900 system, including Airyscan super-resolution detection where indicated. Z-stacks were acquired using 40× water-immersion or 63× oil-immersion objectives. Within an experiment, matched control and POLK-manipulated cultures were acquired using consistent imaging parameters.

Quantitative image analysis was performed with CellProfiler, with ImageJ/Fiji used where indicated. Hoechst or DAPI signals defined neuronal nuclear masks, and MAP2 or TUJ1 immunofluorescence defined neuronal somata and processes. The same segmentation settings, intensity thresholds, size filters, and measurement parameters were applied to corresponding treatment and control images within an experiment. Outcomes included nuclear or somatic median fluorescence intensity, cytoplasmic dsDNA intensity outside the nuclear boundary, signal-positive area, and the number or intensity of individual nuclear 53BP1, γH2AX, PLA, EdU, or other immunofluorescent foci, as appropriate for each assay. Images and individual nuclei or neuronal objects obtained from the same culture well were recorded with their well/chamber identifiers.

### EdU (5-ethynyl-2′-deoxyuridine) assays in human forebrain neurons

Mature human forebrain neurons were incubated with 20 μM EdU for 18 h. This concentration and labeling interval match the conditions used for SAR-seq mapping of endogenous repair-associated DNA synthesis in postmitotic human neurons ^6^.

Cells were washed with PBS. Cells were fixed in 4% paraformaldehyde for 20 min, washed with PBS and permeabilized with 0.5% Triton X-100 for 15 min at room temperature (RT), blocked with 3% BSA in DPBS and subjected to the click reaction for 1.5 h at RT (10 mM copper sulfate, 20 mM THPTA, 20 mg/mL sodium ascorbate, DPBS, and Alexa Fluor azide 594). Cells were washed with DPBS and incubated with 10% goat serum for 1 h.

EdU measurements were restricted to marker-defined neuronal cells or nuclei. EdU intensity was interpreted as neuronal DNA synthesis and, in the context of mature postmitotic cultures and established Repair-seq and SAR-seq studies, as consistent with repair-associated synthesis. EdU incorporation alone does not identify an individual DNA repair pathway.

### EdU Incorporation Assay in primary cortical neurons

EdU incorporation was used to assess DNA synthesis in primary cortical neuronal cultures. Briefly, 5-ethynyl-2′-deoxyuridine (EdU; 10 µM; Thermo Fisher Scientific) was added directly to the culture medium, and cells were incubated for 24 h under standard culture conditions (37°C, 5% CO₂). Following EdU exposure, cells were fixed with 4% PFA and washed with phosphate-buffered saline (PBS). Cells were then permeabilized and processed for EdU detection using the Click-iT™ EdU Cell Proliferation Kit for Imaging (Alexa Fluor™ 594, Cat. No. C10339; Thermo Fisher Scientific), according to the manufacturer’s instructions. Following the Click-iT reaction, cells were blocked and incubated overnight at 4°C with primary antibodies against POLK and MAP2. After washing, cells were incubated with the appropriate fluorescently labelled secondary antibodies for 1h. Nuclei were counterstained with Hoechst, and samples were mounted using mounting medium. Fluorescence images were acquired using a Zeiss LSM confocal microscope.

### SIRF assay

30,000 neurons were plated in 500 µl growth medium in each well of an eight-chamber glass microscope slide (Lab-tek II Chamber Slide 154534) coated with polyornithine, fibronectin and laminin and matured to day 45, with cultures reaching 60-80% confluent on the day of cell labeling. EdU (20 µM) was added 18 h before fixation, and one well was processed without EdU as a negative control. The next day, medium was gently aspirated from the wells and slides were fixed with 4% PFA for 15 min at room temperature (RT). Cells were washed with DPBS. cells were permeabilized with 0.5% Triton X-100 in PBS for 15 min at RT.

The click reaction was performed in a humid chamber equilibrated for 5 min at RT. Fresh Click-iT reaction mix was prepared by combining 2 mM copper sulfate, 10 μM biotin-azide (Invitrogen B10184) and 100 mM sodium ascorbate in DPBS, added in sequence and mixed by vortexing. Sodium ascorbate was freshly prepared, and samples were incubated at RT for 1 h. Blocking was done with 10% goat serum and 0.1% Triton X-100 in PBS at 37°C for 1h.

Primary antibodies against POLK (1:200 SC-166667), MAP2 (1:500, AB-92434), and biotin (1:100; anti-biotin D5A7, Cell Signaling Technology #5597) were diluted in antibody diluent and incubated overnight at 4°C. Wells were washed with Duolink buffer A, and PLA probes (Anti-mouse plus and Anti-rabbit minus Duolink In Situ PLA probe Anti-Rabbit Plus and Anti-Mouse Minus) were added and slides were placed in a prewarmed (37°C) humid chamber.

Wells were washed with Duolink wash buffer A, three times for 5 min each then the ligation reaction was performed with Duolink ligation stock buffer and Duolink ligase enzyme in autoclaved water for 30 min at 37℃ and washed twice with buffer A (Duolink Insitu wash buffer A). Amplification step was performed by diluting Duolink amplification stock and Duolink rolling circle polymerase enzyme in autoclaved water for 100 min at 37℃ and then washed with buffer B (Duolink Insitu wash buffer B 200 mM Tris-HCl pH 7.5 and 100 mM NaCl).

Samples were incubated for 1 h at 37°C with Alexa Fluor 488 goat anti-chicken secondary antibody (Invitrogen A48260) and then mounted with mounting media Duolink In Situ Mounting Medium with DAPI (DUO82040). Confocal imaging used 405, 488, and 561 nm excitation for DAPI, MAP2, and PLA, respectively.

Following EdU incorporation, EdU-containing DNA was conjugated to biotin by copper-catalyzed click chemistry. Proximity between endogenous POLK and biotin-labeled DNA was then detected by PLA using anti-POLK and anti-biotin antibodies. Parallel cultures processed without EdU served as negative controls; both experimental and control samples underwent the same downstream click-labeling, antibody incubation, and PLA procedures. The resulting PLA signal reports molecular proximity, not direct POLK-catalyzed incorporation.

### Proximity Ligation Assay

For protein-proximity analysis in neurons, cells were fixed with 4% paraformaldehyde for 20 min at room temperature, washed three times with PBS and then permeabilized with 0.5%Triton X-100 for 10 min. and washed again with PBS. Incubated for 60 min in blocking buffer provided in the kit at 37℃ in a humidified chamber. Incubated with primary antibodies in antibody diluent overnight at 4℃ and washed with buffer A and then incubated for 1 h at 37℃ in PLA probes (Anti-mouse plus and Anti-rabbit minus Duolink In Situ PLA probe Anti-Rabbit Plus DUO92002 and Anti-Mouse Minus DUO92004), then Duolink In Situ Detection reagents red (DUO92008 Sigma Aldrich) was used, Ligation reaction was performed for 30 min at 37℃ and then again washed with 2 times with buffer A (DUO82049Duolink In Situ wash buffer A). Amplification step was performed for 100 min at 37℃ and then washed 2 times with buffer B (DUO82049 Duolink In Situ wash buffer B). Incubated overnight in primary antibody MAP2 (chicken) and then secondary antibody incubation was done with goat anti-chicken Alexa fluor 647 for 1 h at room temperature and then mounted with mounting media containing DAPI. Confocal microscopy was performed at 405, 561 and 640nm excitation for DAPI, PLA, and MAP2 respectively.

### Senescence-Associated β-Galactosidase (SA-β-gal) Staining

Primary cortical neurons cultured on 8-well chamber slides were rinsed with DPBS+ and fixed with 2% PFA for 10 min at room temperature. Cells were washed twice with 1% bovine serum albumin (BSA) in DPBS+ and incubated with 1× CellEvent™ Senescence Green Working Solution (Invitrogen, Cat. No. C10841) for 2 h at 37°C in a CO₂-free incubator, protected from light. Following incubation, cells were washed three times for 5 min each with 1% BSA in DPBS+. Nuclei were counterstained with Hoechst 33342 (Thermo Fisher Scientific). Samples were mounted using mounting medium, stored at 4°C, and imaged using a Zeiss LSM confocal microscope.

### Sholl Analysis

Dendritic complexity was quantified in MAP2-immunostained primary cortical neurons using Sholl analysis. Individual neuronal dendritic arbors were processed using CellProfiler^54^ (version 4.2.6) and analyzed in ImageJ (NIH) using the Sholl Analysis plugin. The center of each neuronal soma was used as the origin for concentric sampling circles, with an initial radius of 5 µm and subsequent radial increments of 10 µm, extending to a maximum radius of 65 µm. The number of dendritic intersections with each concentric circle was quantified at each radial distance. Sholl profiles were generated by plotting the number of intersections against the distance from the soma. Images were calibrated using the corresponding acquisition-specific pixel dimensions, and identical image-processing and analysis settings were applied to both siControl- and si*Polk*-treated neurons. Data are presented as the mean number of dendritic intersections ± SEM.

### RT-qPCR Analysis in primary cortical neurons

Total RNA was isolated using TRIzol™ Reagent (Thermo Fisher Scientific), according to the manufacturer’s instructions. Complementary DNA (cDNA) was synthesized from 1 µg of total RNA using SuperScript™ IV VILO Master Mix (Invitrogen, Cat. No. 11766050), following the manufacturer’s protocol. Quantitative PCR (qPCR) was performed using PowerUp™ SYBR™ Green Master Mix (Applied Biosystems, Cat. No. A25742) on an Applied Biosystems real-time PCR system. Target gene expression was normalized to the expression of an appropriate internal control gene, and relative gene expression was calculated using the comparative threshold cycle method (2⁻ΔΔCT).

### RNA extraction and qPCR from human forebrain neurons

RNA was extracted from the cells with the Single Cell RNA Purification Kit (Norgen #51800) following the manufacturer’s instructions with some modifications. 200 µl of buffer RL was added to the plates, and lysate was collected; 200 µl of ethanol was added to the lysate and mixed by vortexing for 10 seconds. The lysate/ethanol mixture was applied to a micro spin column onto the column and centrifuged for 1 min at 3,500g. The column was spun for an additional minute at 20,800g and flow-through was discarded, The column was then washed with 400 µl solution A and centrifuged for 1 min at 20,800g flow-through was discarded and wash was repeated and column was spin for 2 min at 20,800g to dry the resin. 15 µl of elution solution A was added and the purified RNA was collected by centrifugation for 1min at 425g followed by 1 min at 20,800g.

The cDNA was prepared from RNA with the iScript cDNA Synthesis kit (Bio-Rad #1708891). Quantitative reverse transcription polymerase chain reaction (qRT-PCR) was performed to measure the level of expression of POLK after 72 h of transfection. Samples from non-targeting control and siPOLK-treated wells were prepared for qRT-PCR. Each sample was run in triplicate on the QuantStudio™ 3 Real-time PCR instrument Applied Biosystems (Thermo Fisher Scientific) with cycling conditions 95°C for 2 min, followed by 40 cycles at 95°C for 15 s and 60°C for 60 s. PowerTrack™ SYBR™ Green Master Mix (# A46109). Each primer was used at a final concentration of 200 nM. Expression levels were normalized with β-actin. The Primers used were as follows.

### Primer sequences used for human neuronal RT-qPCR

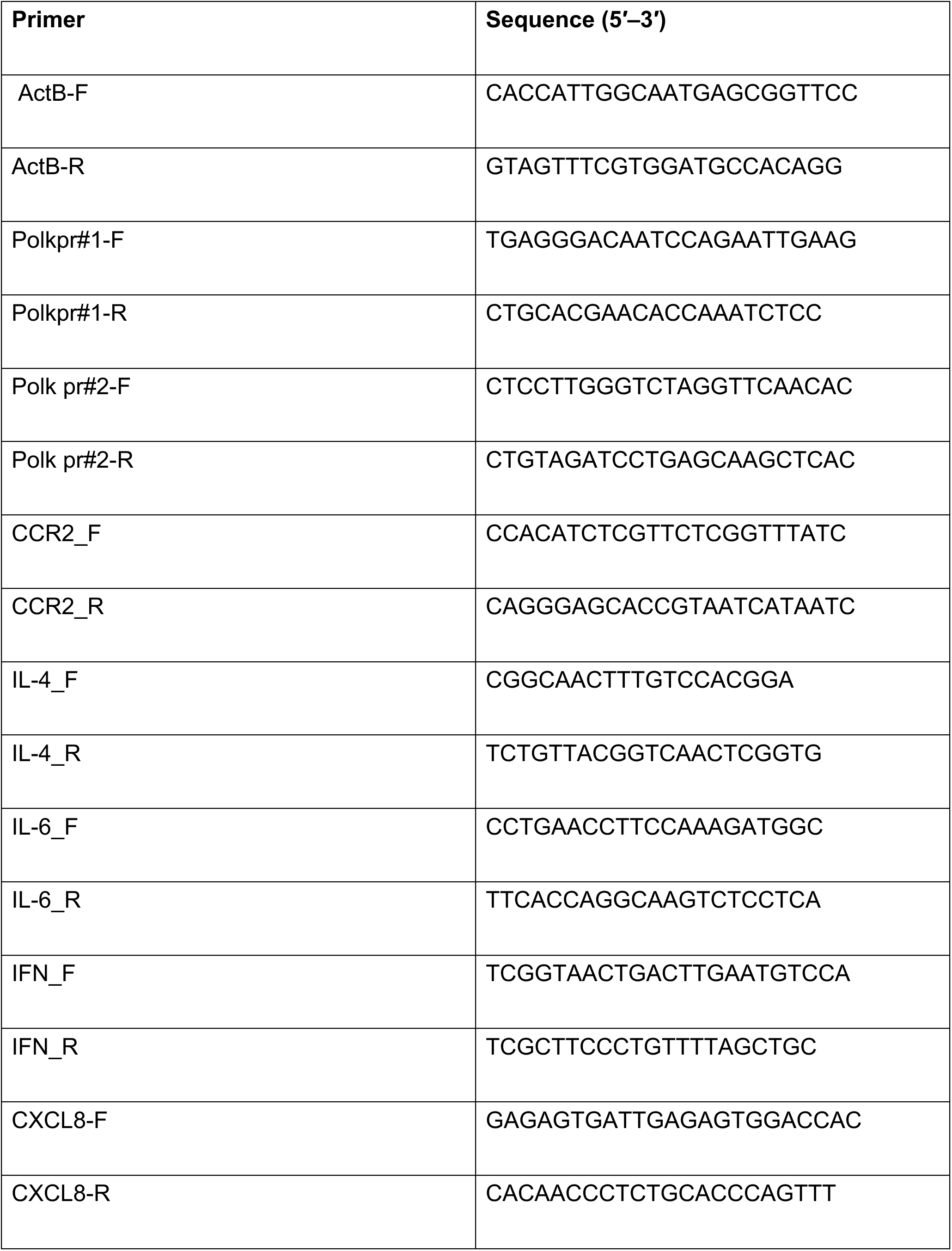
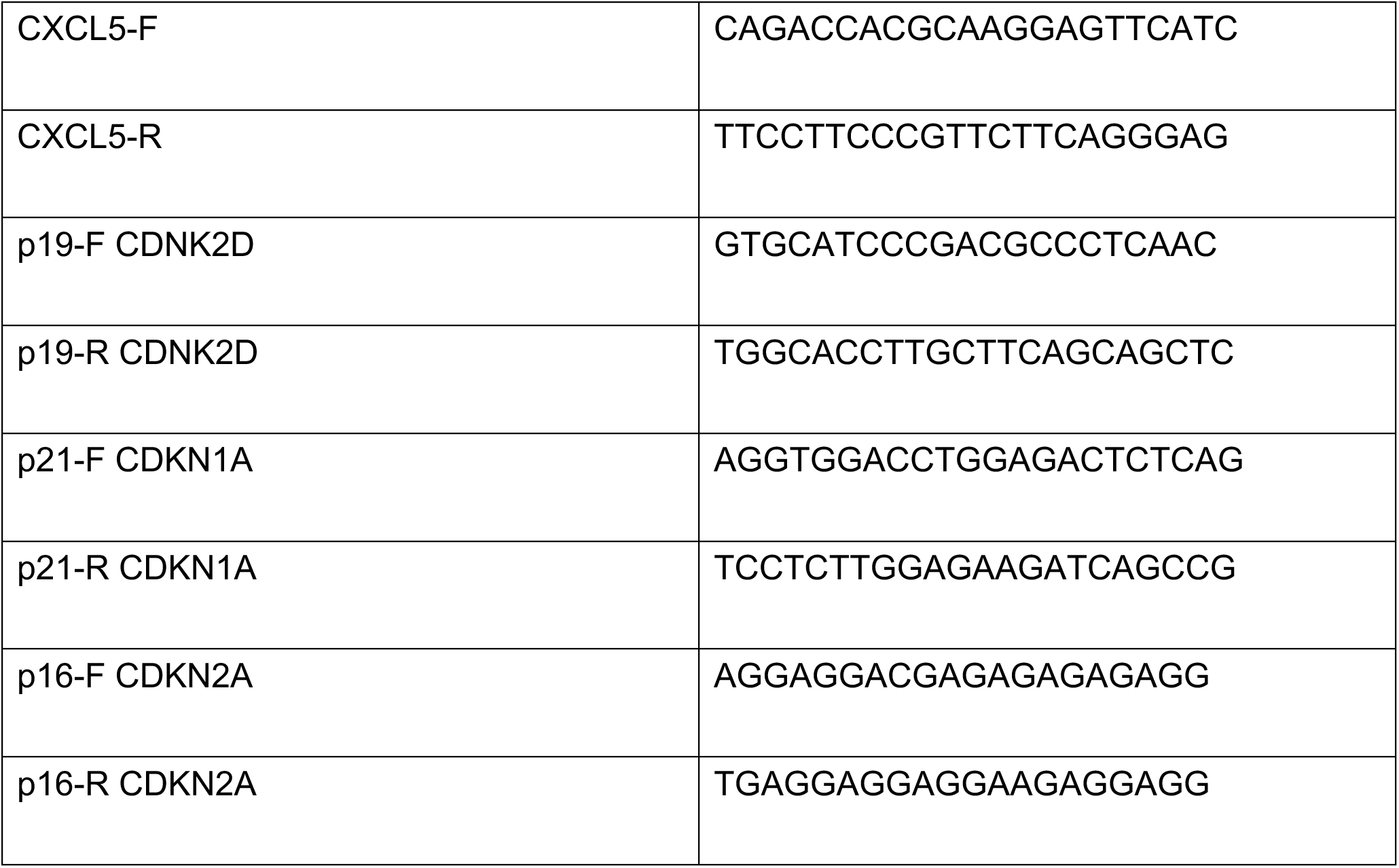

### Protein Extraction and Western Blotting in primary cortical neurons

Total protein was extracted from mouse cerebral cortex tissue and primary neuronal cultures using N-PER™ Neuronal Protein Extraction Reagent supplemented with Halt™ Protease Inhibitor Cocktail (Thermo Scientific), according to the manufacturer’s instructions. Protein concentrations were determined using the Pierce™ BCA Protein Assay Kit (Thermo Scientific). Western blotting was performed as previously described^28^. Briefly, protein samples were denatured, separated by SDS-PAGE using 4–20% Mini-PROTEAN™ TGX™ precast gels (Bio-Rad), and transferred onto polyvinylidene difluoride (PVDF) membranes. Membranes were blocked for 1h in 5% non-fat dry milk and incubated overnight at 4°C with the appropriate primary antibodies. Following washing, membranes were incubated with the corresponding horseradish peroxidase (HRP)-conjugated secondary antibodies. Chemiluminescent signals were developed using SuperSignal™ substrate (Thermo Scientific) and visualized using a ChemiDoc™ Imaging System (Bio-Rad). Band intensities were quantified using ImageJ (NIH) and normalized to the corresponding loading control.

### Western blotting of nuclear and cytoplasmic human neuronal fractions

Nuclear and cytoplasmic samples were prepared with Protease inhibitor cocktail solution (Halt™ Protease Inhibitor Cocktail, EDTA-Free (100X)#18622209) using a nuclear extraction kit (Active motif # 40010). Protein material was quantified using the Pierce™ BCA Protein Assay Kits (#23227). Proteins were resolved in 1× Laemmli reducing buffer by SDS-PAGE electrophoresis and transferred to a nitrocellulose membrane (Bio-Rad). Subsequently, membranes were blocked for 1 h in 5% non-fat milk-1X PBS solution and incubated overnight with primary antibody (Dnase2, 1:200, 15934-1-AP). Membranes were then washed three times in 1X PBS; 0.05% Tween solution and incubated for 1 h with corresponding horseradish peroxidase-conjugated secondary antibody. Membranes were developed using the SuperSignal substrate (Thermofisher # 30496). Blots were quantified using the Image Lab software (Bio-Rad). Antibodies used are listed in supplemental sheet.

### Single-nucleus RNA sequencing and cell-type annotation

Single nuclei were isolated from differentiated human induced pluripotent stem cell (hiPSC) cultures. Single-nucleus gene expression libraries were prepared using the Chromium GEM-X Flex Gene Expression assay (10x Genomics) with four-plex multiplexing, following the manufacturer’s protocol, and sequenced on an Element Biosciences sequencing platform. Raw FASTQ files were processed using Cell Ranger multi (10x Genomics) with the human GRCh38 reference and Chromium Human Transcriptome Probe Set v1.1.0 to generate gene expression count matrices. Log-normalized gene expression profiles were used for cell-type annotation using CellTypist (doi:10.1126/science.abl5197) with an adult human prefrontal cortex reference model (doi:10.1126/science.abo7257). The relative abundance of each neuronal and glial cell type was calculated for each sample as a percentage of the total nuclei assigned to neuronal and glial populations.

### Label-free quantitative proteomics

Human KOLF iPSC-derived forebrain neurons treated with scrambled control siRNA (siControl) or a pool of four POLK-targeting siRNAs (siPOLK; Dharmacon) were collected, washed, and immediately snap-frozen in liquid nitrogen. Cell pellets were thawed on ice and lysed in 100 µL buffer containing 5% SDS, 100 mM triethylammonium bicarbonate (TEAB, pH 8.5), 2× protease inhibitors, and 1× phosphatase inhibitors. Lysates were subjected to probe sonication to shear genomic DNA and clarified by centrifugation at 4°C for 10 min. Protein concentrations were determined using a bicinchoninic acid (BCA) assay (Pierce) according to manufacturer protocol. Proteins were reduced with 3 mM tris(2-carboxyethyl)phosphine (TCEP, 15 min RT), alkylated with 10 mM iodoacetic acid (30 min RT), acidified with phosphoric acid, and diluted fivefold with 90% methanol/100 mM TEAB before loading onto S-Trap Micro columns (ProtiFi). Following three washes with 90% methanol/100 mM TEAB, proteins were proteolyzed overnight with 10 µg trypsin (Sequencing Grade Modified Porcine Trypsin, Promega) at 37°C. Peptides were sequentially eluted using TEAB, 0.2% formic acid, and 50% acetonitrile, dried, and resuspended in loading buffer.

Peptides were separated by nanoscale reversed-phase liquid chromatography using a C18 trap column (10 cm × 100 µm, 5 µm particles) coupled to an analytical C18 column (30 cm × 75 µm, 1.9 µm particles, Dr. Maisch). Separation was performed using a 5–35% acetonitrile gradient in 0.1% aqueous formic acid at a flow rate of 200 nL/min. Eluting peptides were analyzed on an Orbitrap Exploris 480 mass spectrometer (Thermo Fisher Scientific) operated in data-independent acquisition (DIA) mode. Precursor spectra were acquired at 60,000 resolution across m/z 380–985, followed by higher-energy collisional dissociation (HCD; 28% normalized collision energy) and fragment ion acquisition at 15,000 resolution.

Raw mass spectrometry data were analyzed using DIA-NN against a predicted human protein spectral library supplemented with common contaminants (cRAP). Precursor identifications were filtered at a false discovery rate (FDR) below 1%. Label-free quantification was performed by integrating extracted precursor ion chromatogram intensities across samples. A total of 6,732 proteins were identified, of which 5,767 were retained for quantitative analysis after excluding proteins identified by only one peptide. Relative protein abundance was calculated as the log2-transformed intensity ratio between siPOLK and siControl samples. Because the analysis included one sample per condition, protein abundance differences were evaluated descriptively without statistical significance testing. For visualization (Fig. 3), an exploratory pseudo-volcano plot was generated using log2(siPOLK/siControl) on the x-axis and log10-transformed siPOLK protein abundance on the y-axis.

### POLK immunoprecipitation coupled with mass spectrometry (IP-MS)

Human iPSC-derived forebrain neurons were fractionated into nuclear and cytoplasmic fractions, followed by immunoprecipitation using an antibody against DNA polymerase kappa (POLK) or an IgG control. Four immunoprecipitated samples, comprising nuclear POLK-IP, nuclear IgG, cytoplasmic POLK-IP, and cytoplasmic IgG immunoprecipitated protein complexes, were collected on beads in Laemmli buffer and processed for liquid chromatography-tandem mass spectrometry (LC-MS/MS).

Samples were adjusted to a final concentration of 5% SDS in 100 mM triethylammonium bicarbonate (TEAB), followed by reduction with 3 mM tris(2-carboxyethyl)phosphine (TCEP) and alkylation with 10 mM iodoacetic acid. Samples were acidified with phosphoric acid and diluted with 90% methanol/100 mM TEAB before loading onto S-Trap Micro columns (ProtiFi). Following three washes, proteins were digested overnight with 1 µg trypsin at 37°C. Peptides were eluted twice with 80% acetonitrile/100 mM TEAB, dried, and resuspended in 40 µL loading buffer containing 5% dimethyl sulfoxide and 0.1% trifluoroacetic acid. A 4 µL aliquot of each sample was subjected to LC-MS/MS analysis.

Peptides were separated by nanoscale reversed-phase liquid chromatography using a C18 trap column (10 cm × 100 µm, 5 µm particles) coupled to an analytical C18 column (30 cm × 75 µm, 1.9 µm particles), with a 5–35% acetonitrile gradient in 0.1% formic acid at a flow rate of 200 nL/min. Eluting peptides were ionized by electrospray at 2,200 V and analyzed using an Orbitrap Exploris 480 mass spectrometer (Thermo Fisher Scientific) operated in data-independent acquisition (DIA) mode. Precursor ion spectra were acquired at 60,000 resolution across m/z 380–985. Fragmentation was performed using higher-energy collisional dissociation (HCD) at 28% normalized collision energy, and fragment ion spectra were acquired at 15,000 resolution.

Raw mass spectrometry data were analyzed using DIA-NN against a predicted human protein spectral library supplemented with a database of common contaminants (cRAP). Mass tolerances for MS1 and MS2 were determined automatically, and precursor identifications were filtered at a false discovery rate (FDR) below 1%. A total of 6,665 protein groups were identified. Following exclusion of 924 protein groups supported by only one identified peptide, 5,741 protein groups were retained for label-free quantitative analysis. Missing abundance values were imputed using a fixed intensity of 27,000, representing the estimated background noise level. Relative protein enrichment was calculated separately for nuclear and cytoplasmic fractions as the log2-transformed ratio of POLK-IP to the corresponding IgG control. Because the experiment included one sample per condition, the analysis was considered exploratory, and no statistical significance testing was performed.

### Statistical analysis

Statistical analyses and figure preparation were performed using GraphPad Prism 11. For comparisons of two independent experimental groups, the nonparametric Mann–Whitney U test was used; for comparisons involving more than two groups, the Kruskal–Wallis test was used, as appropriate for the experimental design. Unless otherwise indicated, quantitative plots display the mean with 95% confidence intervals (CI). Where reported, the Hodges–Lehmann estimate of the between-group location difference and its 95% CI supplemented the nonparametric hypothesis test.

Most imaging experiments included multiple independently treated neuronal culture wells or chambers per condition, including separate control and POLK-targeting siRNA treatments.

## DATA AVAILABILITY

Mass Spectrometry raw data are available via the MassIVE repository (University of California San Diego) with dataset identifier MSV000103793.manuscript. Additional microscopy images and supporting data are available from the corresponding authors upon reasonable request.

## AUTHOR CONTRIBUTIONS

Gunjan Goyal: hiPSC-derived forebrain-neuron experiments. Premkumar Palanisamy: primary mouse cortical-neuron experiments. Ainesh Paul: hiPSC-derived forebrain-neuron imaging, CellProfiler pipelines, and image quantification. Vijay L. Roy: single nucleus transcriptomics data analysis. Paolo Cifani: Proteomics and analysis. Shilpi Paul and Anirban Paul: conceptualization, resources, supervision, writing - original draft, review, and editing.

## COMPETING INTERESTS

The authors declare no competing interests.

## FUNDING

This work was supported by the National Institutes of Health (RF1AG072602/R01AG072602 to A.P. and 5R03AG081828 to S.P.) and by Startup Research Funds from the Department of Neuroscience and Experimental Therapeutics, Penn State College of Medicine (S.P.).

---

**Supplementary Figure 1.**
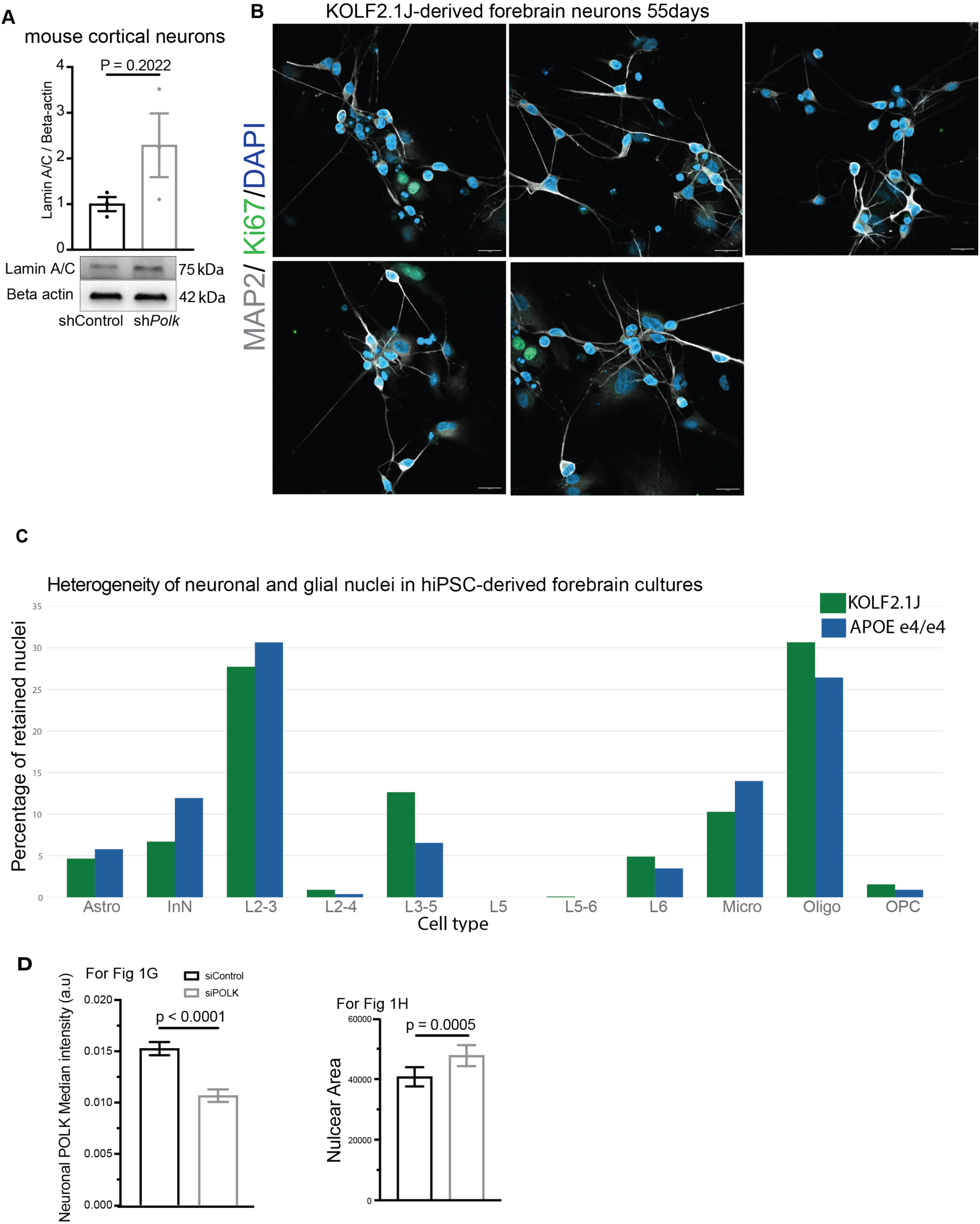
Supporting characterization of POLK-depleted neuronal preparations and human forebrain cultures. (A) Immunoblot for lamin A/C relative to β-actin in mouse cortical neurons after shControl or shPolk; quantification shows a directionally increased but variable signal (P = 0.2022). (B) Representative MAP2, Ki67, and DAPI images of day-55 KOLF2.1J forebrain cultures; Ki67 immunoreactivity is confined largely to MAP2-negative cells. (C) Distribution of annotated neuronal and glial nucleus classes in KOLF2.1J and APOE e4/e4 forebrain cultures based on single-nucleus RNA-sequencing assignments, including astrocytes, interneurons, cortical excitatory neuron classes, microglial-like cells, oligodendroglial cells, and OPCs. The plot represents proportions of retained nuclei in the indicated classes. (D) Supporting quantification of nuclear POLK median intensity (P < 0.0001) and nuclear area (p = 0.0005) for the KOLF2.1J data presented in Fig. 1

**Supplementary Figure 2.**
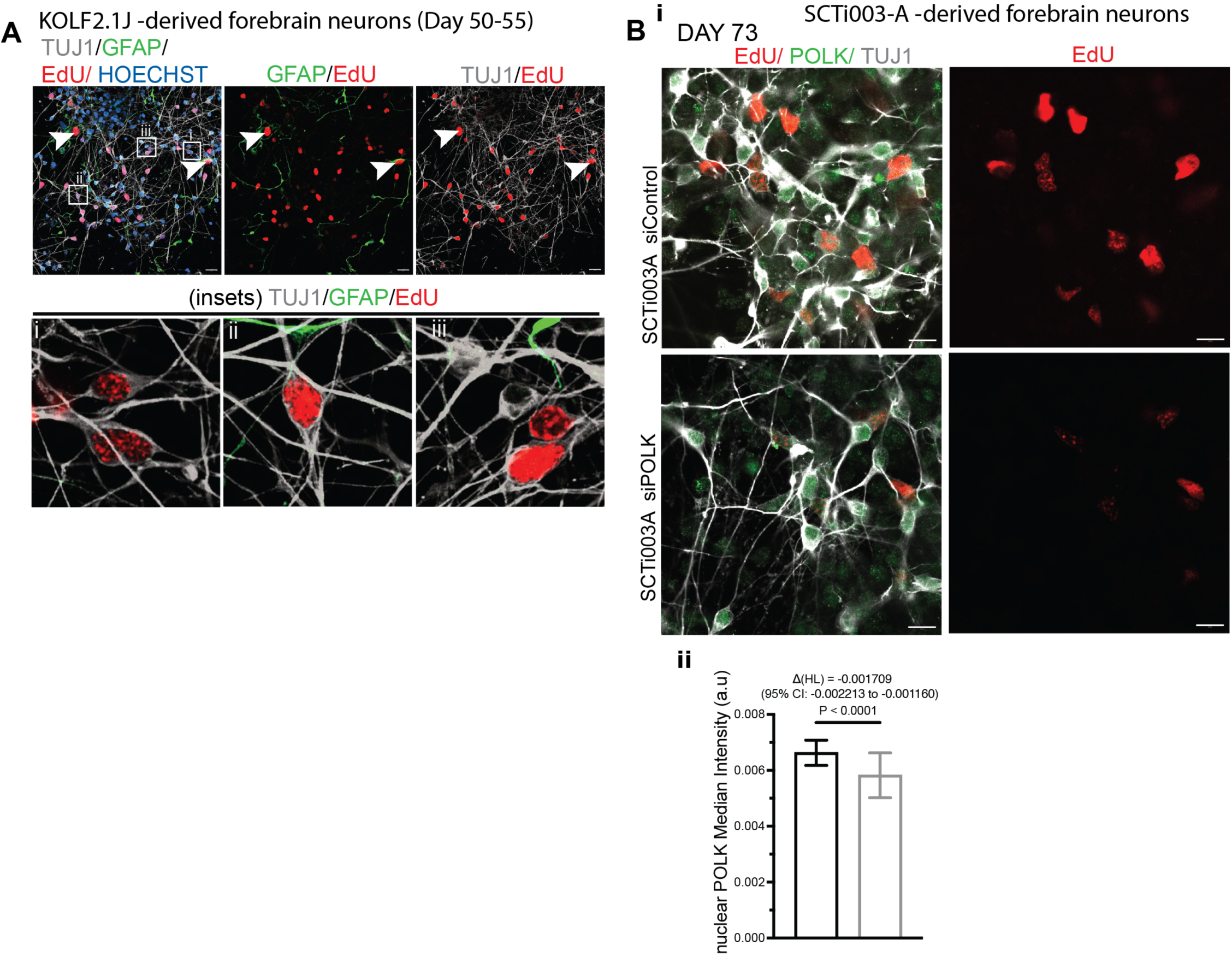
EdU labeling and POLK measurements in independent hiPSC-derived forebrain cultures. (A) KOLF2.1J-derived neurons at days 50–55 stained for TUJ1, GFAP, EdU, and Hoechst, with enlarged examples showing EdU-labeled neuronal nuclei in cultures that also contain GFAP-positive cells. (B) SCTI003-A-derived forebrain neurons at day 73 after siControl or siPOLK treatment. (i) Representative EdU, POLK, and TUJ1 fluorescence images and EdU-only images. (ii) Nuclear POLK median intensity is reduced following POLK depletion (P < 0.0001); the Hodges– Lehmann shift and 95% confidence interval are indicated. These panels support the EdU measurements in Fig 5.

**Supplementary Figure 3.**
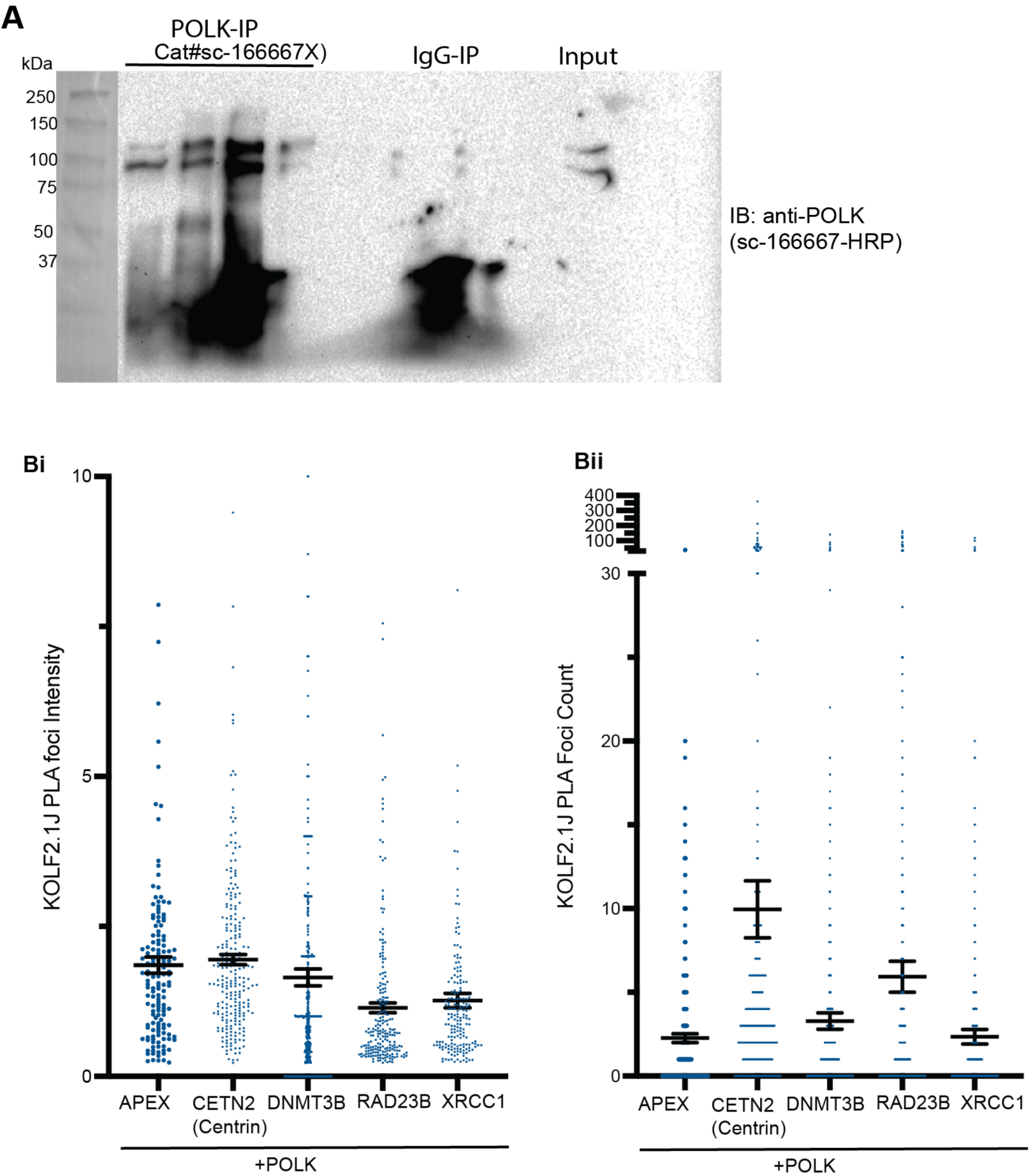
Distribution of KOLF2.1J neuronal PLA signals for additional POLK proximity candidates. (A) POLK-immunoprecipitation samples sent for Proteomics, SDS-PAGE followed by western blot with anti-POLK antibody. (B) (i) Single-cell PLA foci intensity for POLK paired with APEX1, CETN2 (centrin), DNMT3B, RAD23B, or XRCC1. Individual dots represent measured neuronal objects, and horizontal summaries are shown as in the figure. (ii) PLA foci counts for the same candidate pairs. A discontinuous axis accommodates high-count outliers. These plots provide quantitative context for the representative images in Fig. 6C.

**Supplementary Figure 4.**
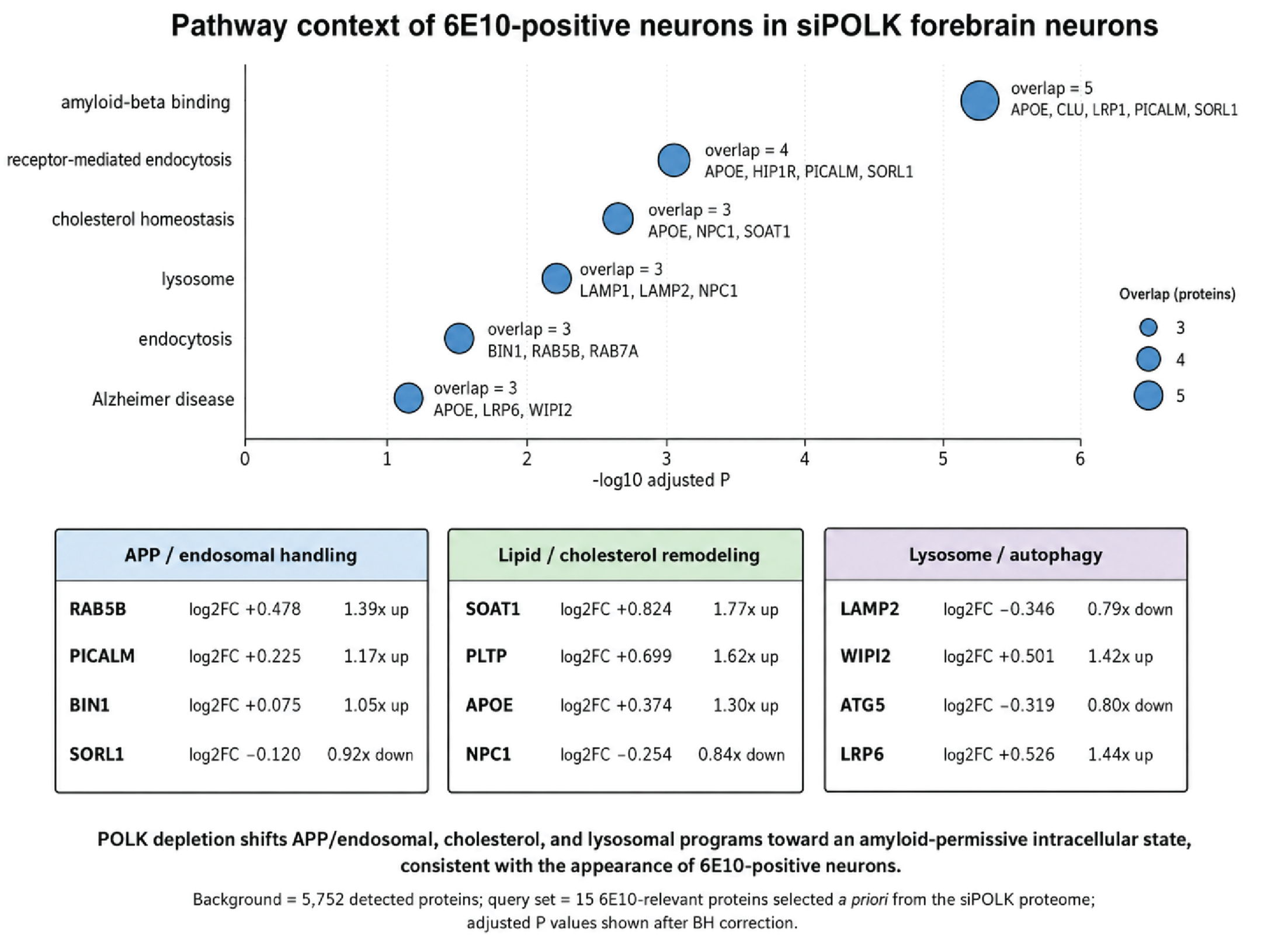
Exploratory APP/endosomal, cholesterol, and lysosomal pathway context for POLK-depleted neurons. Upper panel: Functional enrichment of 15 proteins selected a priori for their relation to 6E10-associated APP/Aβ biology, among 5,752 proteins detected in the exploratory KOLF2.1J siPOLK proteome. Bubble size indicates overlapping protein number, and the horizontal axis shows −log10 of the Benjamini–Hochberg-adjusted enrichment P value; terms include amyloid-beta binding, receptor-mediated endocytosis, cholesterol homeostasis, lysosome, endocytosis, and Alzheimer disease. Lower tables: Descriptive log2 fold changes and corresponding fold changes for selected proteins related to APP/endosomal handling (RAB5B, PICALM, BIN1, SORL1), lipid/cholesterol remodeling (SOAT1, PLTP, APOE, NPC1), and lysosome/autophagy (LAMP2, WIPI2, ATG5, LRP6).

## Notes

### Competing Interest Statement

The authors have declared no competing interest.

### Summary of Updates

Additional and edited figures, updated abstract, updates results and discussion.

